# Genetic investigation of honeybee populations in Kazakhstan

**DOI:** 10.1101/2023.02.08.527617

**Authors:** D. Gritsenko, K. Temirbayeva, A. Taskuzhina, V. Kostuykova, A. Pozharskiy, M. Khusnitdinova, O. Krupskiy, A. Mayer, U. Nuralieva

**Affiliations:** Institute of Plant Biology and Biotechnology, 45 Timiryazev st., 050040, Almaty, Kazakhstan; Kazakh Scientific Research Institute of Animal Husbandry and Forage Production, 51 Zhandosov str., 050035, Almaty, Kazakhstan; al-Farabi Kazakh National University, 71 al-Farabi Ave., 050040, Almaty, Kazakhstan

**Author notes:** **Correspondence**: D. Gritsenko –.

**Keywords:** Honeybee, population, haplotype, SSR, cluster analysis, genetic distance, cross-breeding, genetic admixture, diversity

## Abstract

Beekeeping as a staple of agriculture in Kazakhstan is believed to have emerged when external bee specimens were introduced into the country. The Central Russian bee (*Apis mellifera mellifera*) has been present throughout Kazakhstanian apiaries for a long time since its import into Eastern Kazakhstan at the end of the 18^th^ century. To date, six subspecies have been distributed across the country (*A.m. sossimai*, *A.m. carpatica*, *A.m. mellifera*, *A.m. ligustica, А.m. caucasica*, and *A.m. carnica*). According to mitochondrial haplotype analysis based on DraI mtDNA COI-COII (DmCC) test, local samples were represented by C2 (316), C1 (99), and M4 (7) haplotypes. The results of simple sequence repeats (SSR) genotyping revealed a large polymorphism at nine microsatellite loci, with the number of alleles amounting to 35 (AP55), 32 (AP43), 25 (A124), 18 (A113), 13 (A88), 12 (A43), 11 (A007), 7 (A28), and 5 (A24). Relative to the expected heterozygosity (H_e_), the observed heterozygosity (H_o_) was slightly higher for most markers considering both the overall samples and individual populations. The inbreeding coefficient confirmed the excess outbreeding for the geographical populations of car-Shym-T (−0,105), car-zham-A (−0,114), Zhet-Alakol (−0,008), and Zhet-Ushbulak (−0,028). The admixture of honeybee local populations was confirmed by the research presented here.The differentiation of populations was only possible by geographical location according to clustering analysis. A considerable degree of genetic admixtures among subspecies was identified in every population. The subspecies were not separated from each other. None of the groups formed by the neighbor-joining tree based on Nei’s genetic distance included a precise subspecies or population. All groups represented an admixture of subspecies from different populations.

Unregulated cross-breeding for the past 50 years has laid the foundation for the promiscuous genetic nature of honeybee populations in Kazakhstan. It could be concluded that some samples were the result of cross-breeding with endemic bee *Apis mellifera pomonella* since most apiaries were located in areas of endemic bee distribution.

## 1. Introduction

*Apis mellifera* L. or the Western honeybee is a species of social bees belonging to the *Apidae* family. The *Apinae* subfamily is widely distributed on all continents except Antarctica. *A. mellifera* L. can be divided into five evolutionary lineages (A, M, C, O, Y, and Z) and consists of 27-30 subspecies according to morphological and phylogeographic research. This large number of subspecies was formed due to the isolation and subsequent accumulation of genetic differences. The intensive hybridization of the species is ongoing [1–4].

Several hypotheses based on morphological and DNA analysis data have been proposed to explain the origins of the subspecies and diversity of evolutionary lines. Asia, the Middle East, and Africa are considered possible centers for the origin of *A. mellifera* L., with subsequent bee migration to Eurasia via the Iberian and Arabian peninsulas [1,2,4,5].

The territory of Kazakhstan is vast and covers various types of natural conditions, including those allowing for the development of beekeeping. Archeological studies of historical monuments in Altai (Kazakhstan side) and Siberia, as well as research into the history of Altai languages suggest that bees inhabited the territory of present-day Altai in ancient times. Beekeeping as a staple of agriculture in the region is thought to have emerged when external bee specimens were introduced into the area. The role of local bees in this process has received relatively little research attention until now.

The history of beekeeping in Kazakhstan dates back over more than 200 years to the first mention of the import of bees to the Ust-Kamenogorsk fortress. According to available information [4], bees were first brought to the Ust-Kamenogorsk region in 1786 and then spread across the Altai Mountains and Siberia. The bees had been imported into Kazakhstan from several locations including Bashkiria, Ukraine, and Orenburg [6]. However, according to their phenotype, those bees had likely descended from the European dark bee.

The development of beekeeping in Kazakhstan quickly skipped to the apiary (beehive beekeeping) stage, bypassing the first two customary periods - bee hunting and wild beekeeping. Nevertheless, Sheppard and Meixner [7] consider the honeybee subspecies Apis mellifera pomonella to be endemic to Kazakhstan. This subspecies is believed the result of the crossbreeding between *Apis mellifera* and *Apis cerana* [8].

The main bee subspecies bred in Kazakhstan for commercial use include *A.m. sossimai*, *A.m. carpatica*, *A.m. mellifera*, and *A.m. carnica*. The primary scope of this research is to reveal the genetic structures of honeybee populations in Kazakhstan, their subspecies-based distribution, and identify a genetically pure pool for further breeding.

## 2. Material and Methods

### 2.1 Sampling and DNA isolation

Specimens of *Apis mellifera* L. were collected from 36 populations during the summer-autumn periods of 2021 and 2022 on the territory of the Republic of Kazakhstan. These included individuals from the *A.m. carpatica, A.m. sossimai, А.m. caucasica, A.m. ligustica*, and *A.m. carnica* subspecies. The *Apis mellifera mellifera* sample was obtained from the Belukha mountain (Russian Federation). Fifteen samples of *A.m. carnica* were provided by Prof. Dr. Kaspar Bienefeld from the Institute for Bee Research Hohen Neuendorf and Humboldt University of Berlin and marked as belonging to the C_group. The location of origin for each population is provided in Table 1, Figure 1.

**Table 1.**
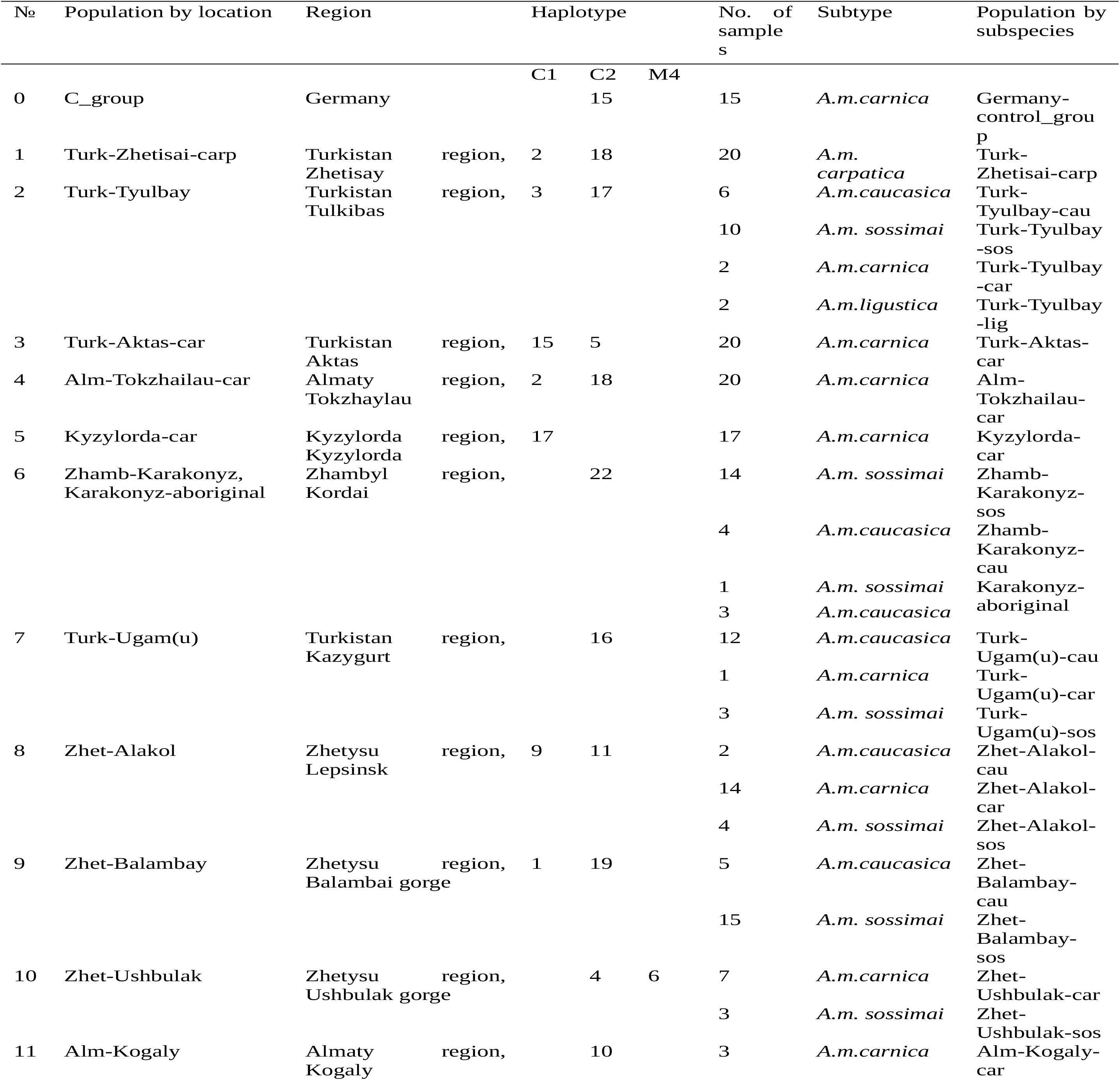
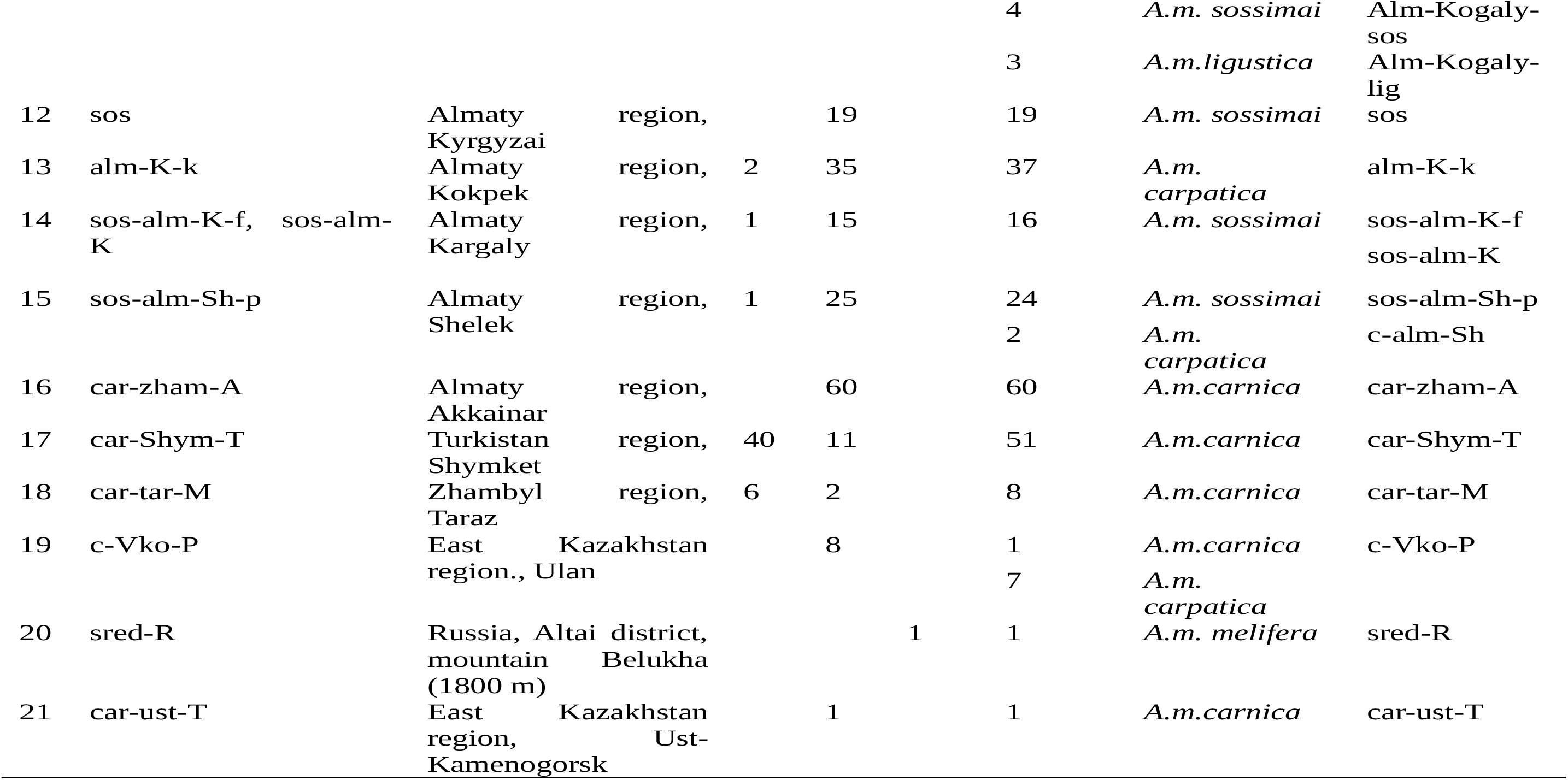
Bee population characteristics

**Figure 1.**
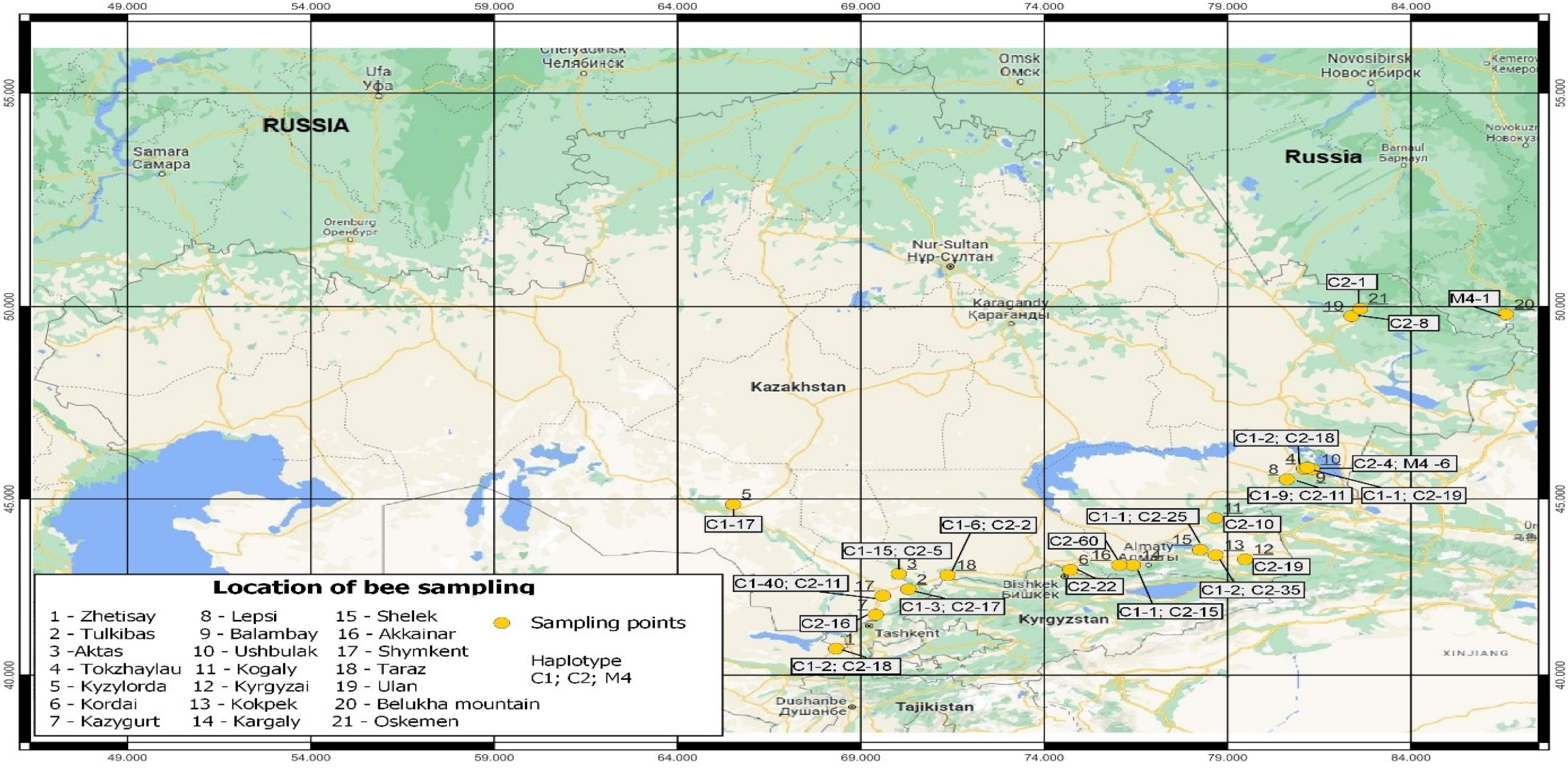
The map of bee sampling locations in Kazakhstan

DNA was isolated from a worker bee from every potential colony following a modified CTAB (cetyltrimethylammonium bromide) method [9]. The thorax or legs of every bee were homogenized in liquid nitrogen with further incubation in 1 mL of a CTAB buffer (4% CTAB, 100 mM tris (pH 8.0), 1.4 M NaCl, 20 mM ethylenediaminetetraacetic acid (pH 8.0), and 0.4% β-mercantoethanol added before use). The incubation lasted for one hour in a thermo-shaker at 65°C with a shaking frequency of 1,000 rpm. The separation of a water phase containing nucleic acids from the organic contents was performed by adding the same volume of a chloroform reagent to homogenized samples followed by 5-7 inverting times of the tubes. After centrifugation at 13,000 rpm for 15 min, the water phase was transferred to a new tube containing 500 ml of cold isopropanol. During this step, the samples were left to rest at 4°C overnight. Following the precipitation of genomic DNA over 15 min at 13,000 rpm, the DNA pellets were washed with 75% ethanol. The dried pellets were then dissolved in 100 uL of ultra-pure water. The DNA concentration was assayed by spectrophotometry (Nanodrop One) and diluted to 10 ng/uL for PCR. The DNA stocks were preserved at −80°C for long-term storage.

### 2.2 mtDNA analysis

All samples (n=437) were analyzed for the COI-COII marker. The PCR amplification of the intergenic COI-COII region was performed using the E2 (5′-GGCAGAATAAGTGCATTG −3′) and H2 (5′-CAATATCATTGATGACC −3′) primers [10]. The size of the amplified DNA fragments was determined by electrophoresis of the PCR product in 1.5% agarose gel. To identify the haplotypes, the PCR products were digested by *DraI* and evaluated on a 10% polyacrylamide gel. The DraI mtDNA COI-COII (DmCC) test was performed according to the protocol described by Garnery et al. [10]. The sequencing of the PCR products obtained by the E2 and H2 primers was performed for each new haplotype (haplotypes having a unique band pattern). PCR was carried out by Phusion Hot Start II High-Fidelity DNA polymerase (Thermo fisher Scientific) followed by the purification of the PCR products with the GeneJET PCR Purification Kit (Thermo fisher Scientific). The BigDye™ Terminator v3.1 Cycle Sequencing Kit (Thermo fisher Scientific) was applied during the sequencing of PCR products conducted on an Applied Biosystems 3500 Genetic Analyzer according to the manufucturer’s protocol. Then, the obtained sequences were aligned with *Apis mellifera* reference sequences [11].

### 2.3 Microsatellite analysis

All honeybee individuals were genotyped by nine microsatellite loci (A113, A43, A28, A24, A88, A007, Ap55, Ap43, A124) [12,13]. Fragment analysis was performed on a 3500 Genetic Analyzer (Applied Biosystems, California, USA). Nine SSR markers were grouped into two multiplexes considering size and fluorescent dye. The first plex included markers A24, A28, A43, A88, A124; the second one contained the remaining markers. Every marker was amplified in 20 μl of a reaction mix containing a 1X Standart Taq Reaction Buffer (M0273E, New England BioLabs Inc.), 0.4 μl of 10 mM dNTP, 0.4μl of 10 mM forward and reverse primers, and 1U Taq-polymerase (M0273E, New England BioLabs Inc.); the mix was adjusted to 20 μl using ddH_2_O. The amplification of the first plex markers and markers A007 and A113 was performed according to the program described in Dunin et al. [14]. Markers Ap55 and Ap43 were amplified following the protocol described in Delaney et al. [12]. Amplicon verification was performed by gel electrophoresis using 1.5% agarose gels and the PCR products were visualized using a Gel Doc XR+ Imaging System (Bio-Rad Inc.). The PCR products were then purified with the GeneJET PCR Purification Kit (Thermo fisher scientific) and diluted to 30 ng/uL. Each plex contained the reaction mix including 1 μl of purified PCR product for each marker, 0.15 μl of GeneScan 500 LIZ Size Standard (Applied Biosystems), 8.85 μl of HI-DI Formamide (Applied Biosystems), and the reaction mix was adjusted to 50 μl using ddH_2_O. After denaturation for 4 min at 95°C, the mix was subjected to capillary electrophoresis using the FragAnalysis50_POP7 manufacture module.

### 2.4 Statistical and phylogenetic analysis

The genotypes were determined based on their fluorescence peaks in the GeneMapper 6 software (Applied Biosystems). The population structure was analyzed using the STRUCTURE2.3.4 software [15] with the following settings: admixture model with no linkage, prior use of population information, 20,000 burn in, and 40,000 MCMC iterations. The analysis was run for K from 1 to 10 with ten repeats. The CLUMPAK web server [16] was used to visualize the results and determine the optimal K value according to Evanno’s method [17]. Phylogenetic analysis was conducted and populations’ genetical statistics were calculated using the ‘adegenet’ [18], ‘poppr’ [19] and ‘ape’ [20] packages in R4.2.1. The distance matrix was calculated using Nei’s genetic distance and then used to build a neighbor-joining tree. The tree was visualized in the FigTree software [21].

## 3. Results

### 3.1 Sampling

The samples of bees were collected from the most promising apiaries located predominantly in Southern and Southeastern Kazakhstan since these regions are characterized by favorable climatic conditions and a flowering plant diversity for beekeeping (Figure 1). Very few bee colonies have been registered in the northern regions, and beekeeping is poorly developed in the west of the country. The collected samples belonged to well-known subspecies, namely, *A.m. carpatica, А.m. caucasica, A.m. ligustica, A.m. carnica, and A.m. mellifera*, as well as to an unconfirmed *А.m. sossimai* one (Table 1). *А.m. sossimai* is broadly distributed in Kazakhstanian apiaries and originated in Ukraine [22]. Half of the apiaries comprised more than one bee subspecies. The Tulkibas apiary in Turkistan included the families of *A.m. carpatica, А.m. caucasica, A.m. ligustica, A.m. carnica*, and *А.m. sossimai*. Every subspecies except for *A.m. ligustica* was also presented separately from the others. The average age of the apiaries was about five years and their genealogical data had been described in some cases. Most of the apiaries were established by bee queens from European lines of *Apis mellifera* L.. The apiaries from the Karakonyz gorge, Zhambyl region, and Ugam gorge in Turkistan were established 50 years ago. The apiary from the village of Tokzhailau in the Almaty region is around 25 years old (Table 1). The sample of *A.m. mellifera* and 17 samples from Germany were used as reference groups in the genetic analysis. Indeed, the lines of honeybees from Germany and Ukraine served as primary material in the establishment apiaries in Kazakhstan. These reference groups were used to identify hydrids and understand the genetic diversity among local populations.

The samples were divided according to their geographical location and the subspecies of every location used in the current research. A total of 22 populations grouped by geographical location and 36 subspecies-based populations were analyzed (Table 1, Figure 1).

### 3.2 Diversity of mitochondrial haplotypes

The DNA amplification and restriction of the COI-COII intergenic region were successful for all analyzed individuals (n=437). The analysis of the COI–COII gene region revealed the presence of European lineages C and M within the populations. Haplotypes C1, C2, and M4 were identified in this study according to the DmCC test (Figure 2). The families in the apiaries mostly bore the haplotypes C1 and C2. The M4 haplotype was identified in six out of ten families from the Ushbulak apiary, Zhetysu region, since a native queen bee was used in their breeding process. The sample from the Belukha mountain also presented an M4 lineage. The PCR products of two or four samples for every haplotype were sequenced using E2 and H2 primers. The C1 haplotype included 420, 64, 47, and 41 bp bands, while the C2 haplotypes contained 420, 64, 47, and 40 bp bands according to the sequencing analysis (Figure 2 and Figure S1). These haplotypes have also been previously described in several studies [23,24]. Twelve of the studied populations included families of both C1 and C2 haplotypes, while ten populations were represented only by the C2 haplotype together with the C_group and only families from the Kyzylorda population included the C1 haplotype. Overall, 338 samples were represented by the C2 haplotype, 99 samples by C1 the haplotype, and seven by the M4 haplotype. The apiary from the Kyzylorda region included 17 families with a C1 haplotype. It was established 32 years ago by the bee queen of Peshetz line 33 – 00033 and breeding within it has since continued without the addition of new lineages or haplotypes. The moststudied populations are being formed by constant introducing bee queens of new subspecies or breeds. The Ushbulak apiary included families of lineages M and C, the apiary was established by a native queen. M4 haplotypes were identified in two families of *А.m. sossimai* and four families of *А.m. carnica*. Most subspecies-diverse apiary, Tulkibas, was represented by three families of C1 haplotype and 17 families of C2 haplotype. With the exception of the apiary from Kyzylorda, the apiaries including only one bee subspecies also consisted of haplotypes C1 and C2.

**Figure 2.**
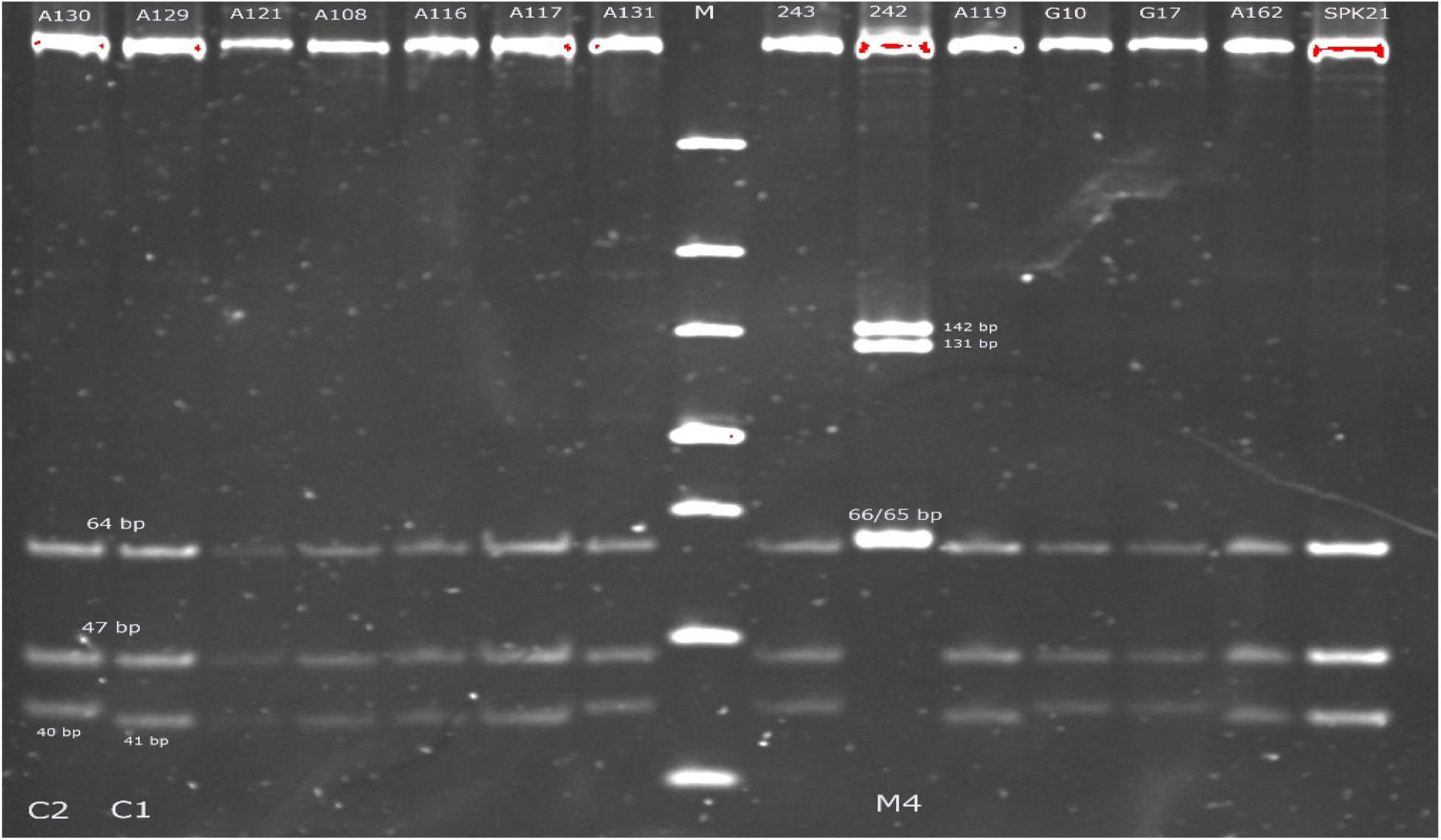
Haplotype differentiation by DraI mtDNA COI-COII test. Haplotypes C1 (Q sequence - 420, 64, 47, 40 bp), C2 (Q sequence - 420, 64, 47, 41 bp), and M4 (PQQ sequence – 422, 142, 131, 66, 65 bp) identified among samples of populations Zhet-Alakol (A group samples), Kyzylorda-car (G10, G17), Turk-Aktas-car (SPK21), car-ust-T (243), and sred-R-Be (242). M – Ultra low range DNA ladder (Invitrogen). The DNA restriction fragments were analyzed by electrophoresis in 10% polyacrylamide gel.

### 3.3 Genetic variability of microsatellite loci

The genetic diversity of 36 bee populations distributed on the territory of Kazakhstan were analyzed using nine informative SSR markers. SSR genotyping raw data is available in Table S1. A total of 437 samples, including 15 samples from a control group, were represented in a descending order of abundance by *A.m. carnica* (n=205), *А.m. sossimai* (n=115),*A.m. carpatica* (n=64), *А.m. caucasica* (n=32), *A.m. ligustica* (n=5), and *А.m. mellifera* (n=1). Polymorphism at nine microsatellite loci varied greatly, with the number of alleles amounting to 35 (AP55), 32 (AP43), 25 (A124), 18 (A113), 13 (A88), 12 (A43), 11 (A007), 7 (A28), and 5 (A24) (Table 2). All markers were polymorphic, and 158 alleles were revealed across the nine loci among 36 populations. The maximum number of alleles across all loci was identified in the Turk-Ugam(u)-cau and sos (68) while the minimum diversity of alleles was found for the Germany control group (42) considering populations of at least ten samples divided by subspecies and location. The observed heterozygosity (H_o_) across all samples ranged from 0.289 (A28) to 0.909 (A124). The highest H_o_ was observed in the Zhet-Alakol-car population across *A.m. carnica* for markers A124 (1.0) and AP43 (1.0), and in the Turk-Ugam(u)-cau *А.m. caucasica* species for marker A124 (1.0). Marker A28 was represented in alm-K-k with the lowest value of H_o_ (0.02). The H_o_ was slightly higher than the H_e_ for most markers considering both the overall samples and individual populations. The mean allelic richness ranged from 1.22 for Turk-Ugam(p)-car *A.m. carnica* to 1.75 for Turk-Ugam(u)-cau *А.m. caucasica*, while a richness value of 1.55 was identified for the control group (Table 3). Identical genotypes were not found in the samples.

**Table 2.** Number of alleles (N_allele), minor allele frequency (MAF), expected (He) and observed (Ho) heterozygosity per marker across samples.

| Marker | N_allele | MAF | He | Ho |
| --- | --- | --- | --- | --- |
| A007 | 11 | 0,225917 | 0,582569 | 0,679401 |
| A113 | 18 | 0,114679 | 0,506881 | 0,60961 |
| A124 | 25 | 0,126147 | 0,81422 | 0,909924 |
| A24 | 5 | 0,413991 | 0,637615 | 0,6205 |
| A28 | 7 | 0,112385 | 0,272936 | 0,289235 |
| A43 | 12 | 0,12156 | 0,538991 | 0,583684 |
| A88 | 13 | 0,181193 | 0,575688 | 0,642404 |
| AP43 | 32 | 0,269495 | 0,699541 | 0,784196 |
| AP55 | 35 | 0,145642 | 0,752294 | 0,870418 |

**Table 3.**
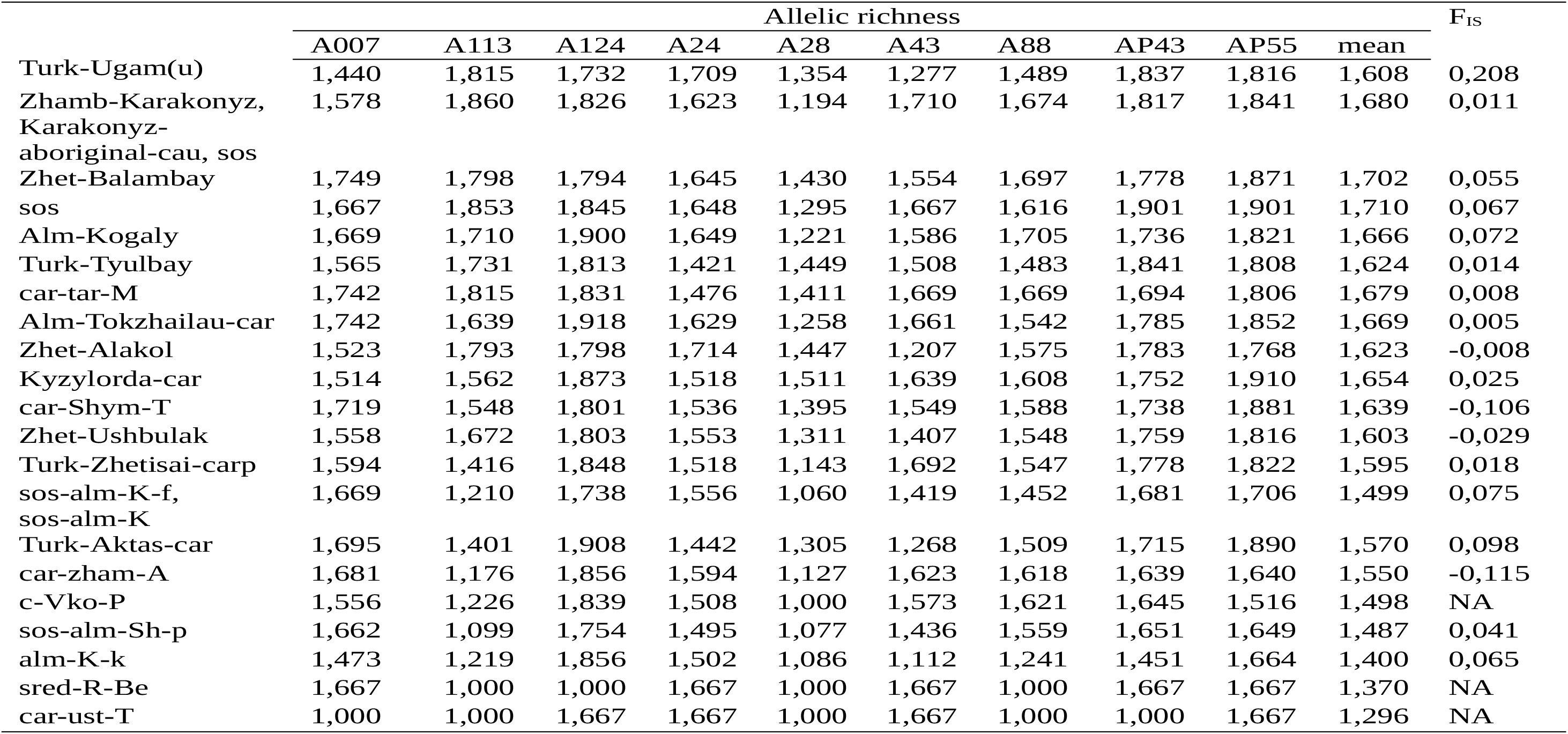
Allelic richness per marker and inbreeding coefficient (F_IS_) among geographical populations

The maximum inbreeding coefficient (*F_IS_*) (0.208) was identified in Turk-Ugam(u) across *А.m. caucasica* comprising fewer heterozygotes than expected (Table 3). Seven out of nine markers showed lower H_o_ than H_e_, while only markers A124 and A113 indicated the opposite trend. The H_o_ for markers A28 and A43 in the Turk-Ugam(u)-cau population were two times lower than the H_e._ Most populations tended to be consistent with the Hardy-Weinberg equilibrium according to *F_IS_* values which were lower than 0.098. Negative values of *F_IS_* were observed for the car-Shym-T (−0,105), car-zham-A (−0,114), Zhet-Alakol (−0,008), and Zhet-Ushbulak (−0,028) geographical populations which comprised more heterozygotes than expected, thus confirming the excess outbreeding.

### 3.4 Genetic relationships among populations

Considering the genetic diversity among the geographically divided populations, the maximum genetic distance (0.502) was found between Zhamb-Karakonyz and the control C_group while the minimum value (0.044) was identified between the sos-alm-Sh-p and alm-K-k according to the Nei’s algorithm (Table S2). The maximum genetic distance (0.516) was found for the subspecies-based populations between *А.m. sossimai* of Zhamb-Karakonyz-sos and the Germany control group; the minimum distance (0.044) was identified between the same populations as the geographically assigned populations. The fixation index (*F_ST_*) values ranged from 0.011 (Zhet-Balambay/Turk-Tyulbay) to 0.108 (Zhamb-Karakonyz, aboriginal/ alm-K-k) (Table 4). The populations containing less than ten samples were not considered.

**Table 4.** Pairwise F_ST_ values between 21 populations of honeybee distributed in Kazakhstan. The F_ST_ values for pairwise comparisons among populations calculated from SSR data. Significant tests were performed using 1,000 permutations. Population name in bold comprised less than ten samples. Bold numbers indicate the maximum and minimum values.

|  | alm-K-k | **Alm-Kogaly** | Alm-Tokzh | **c-Vko-P** | car-Shym-T | **car-tar-M** | **car-ust-T** | car-zham-A | C_group | Kyzylorda | **sred-R-Be** | Turk-Aktas | Turk-Tyulbay | Turk-Ugam | Turk-Zhetisai-carp | sos | sos-alm | sos-alm-Sh-p | Zhamb-Karak | Zhet-Alakol | Zhet-Bal | **Zhet-Ushbulak** |
| --- | --- | --- | --- | --- | --- | --- | --- | --- | --- | --- | --- | --- | --- | --- | --- | --- | --- | --- | --- | --- | --- | --- |
| alm-K-k | - |  |  |  |  |  |  |  |  |  |  |  |  |  |  |  |  |  |  |  |  |  |
| **Alm-Kogaly** | 0,059 | - |  |  |  |  |  |  |  |  |  |  |  |  |  |  |  |  |  |  |  |  |
| Alm-Tokzh | 0,037 | 0,018 | - |  |  |  |  |  |  |  |  |  |  |  |  |  |  |  |  |  |  |  |
| **c-Vko-P** | 0,020 | 0,041 | 0,017 | - |  |  |  |  |  |  |  |  |  |  |  |  |  |  |  |  |  |  |
| car-Shym-T | 0,080 | 0,044 | 0,037 | 0,062 | - |  |  |  |  |  |  |  |  |  |  |  |  |  |  |  |  |  |
| **car-tar-M** | 0,052 | 0,021 | 0,001 | 0,017 | 0,029 | - |  |  |  |  |  |  |  |  |  |  |  |  |  |  |  |  |
| **car-ust-T** | 0,092 | 0,005 | -0,013 | 0,022 | 0,035 | -0,009 | - |  |  |  |  |  |  |  |  |  |  |  |  |  |  |  |
| car-zham-A | 0,050 | 0,052 | 0,030 | 0,038 | 0,061 | 0,038 | 0,046 | - |  |  |  |  |  |  |  |  |  |  |  |  |  |  |
| C_group | 0,047 | 0,048 | 0,020 | 0,047 | 0,063 | 0,016 | 0,042 | 0,026 | - |  |  |  |  |  |  |  |  |  |  |  |  |  |
| Kyzylorda | 0,049 | 0,028 | 0,007 | 0,023 | 0,034 | 0,005 | 0,025 | 0,040 | 0,038 | - |  |  |  |  |  |  |  |  |  |  |  |  |
| **sred-R-Be** | 0,218 | 0,028 | 0,069 | 0,168 | 0,114 | 0,052 | NA | 0,160 | 0,122 | 0,087 | - |  |  |  |  |  |  |  |  |  |  |  |
| Turk-Aktas | 0,040 | 0,035 | 0,019 | 0,041 | 0,037 | 0,026 | 0,048 | 0,057 | 0,043 | 0,024 | 0,083 | - |  |  |  |  |  |  |  |  |  |  |
| Turk-Tyulbay | 0,086 | 0,032 | 0,036 | 0,060 | 0,045 | 0,021 | 0,043 | 0,077 | 0,066 | 0,041 | 0,047 | 0,054 | - |  |  |  |  |  |  |  |  |  |
| Turk-Ugam | 0,082 | 0,032 | 0,041 | 0,065 | 0,060 | 0,033 | 0,007 | 0,085 | 0,072 | 0,038 | 0,048 | 0,056 | 0,019 | - |  |  |  |  |  |  |  |  |
| Turk-Zhetisai | 0,026 | 0,017 | 0,014 | 0,024 | 0,047 | 0,010 | 0,000 | 0,031 | 0,018 | 0,021 | 0,097 | 0,022 | 0,053 | 0,051 | - |  |  |  |  |  |  |  |
| sos | 0,063 | 0,017 | 0,022 | 0,036 | 0,036 | 0,015 | -0,029 | 0,052 | 0,048 | 0,025 | 0,061 | 0,045 | 0,028 | 0,014 | 0,034 | - |  |  |  |  |  |  |
| sos-alm | 0,030 | 0,027 | 0,012 | 0,017 | 0,056 | 0,020 | -0,004 | 0,030 | 0,019 | 0,031 | 0,125 | 0,042 | 0,066 | 0,067 | 0,008 | 0,043 | - |  |  |  |  |  |
| sos-alm-Sh-p | 0,015 | 0,050 | 0,031 | 0,023 | 0,073 | 0,046 | 0,074 | 0,044 | 0,053 | 0,038 | 0,181 | 0,041 | 0,068 | 0,073 | 0,027 | 0,058 | 0,023 | - |  |  |  |  |
| Zhamb-Karak | **0,109** | 0,031 | 0,042 | 0,071 | 0,067 | 0,037 | 0,036 | 0,094 | 0,082 | 0,050 | 0,060 | 0,072 | 0,027 | 0,020 | 0,066 | 0,022 | 0,074 | 0,095 | - |  |  |  |
| Zhet-Alakol | 0,044 | 0,025 | 0,010 | 0,024 | 0,033 | 0,011 | 0,000 | 0,047 | 0,041 | 0,014 | 0,080 | 0,024 | 0,025 | 0,028 | 0,022 | 0,013 | 0,042 | 0,040 | 0,047 | - |  |  |
| Zhet-Bal | 0,076 | 0,019 | 0,014 | 0,046 | 0,039 | 0,016 | 0,005 | 0,061 | 0,054 | 0,021 | 0,060 | 0,039 | **0,012** | 0,016 | 0,037 | 0,012 | 0,053 | 0,057 | 0,018 | 0,010 | - |  |
| **Zhet-Ushbulak** | 0,054 | 0,021 | 0,011 | 0,033 | 0,041 | 0,013 | 0,006 | 0,057 | 0,042 | 0,015 | 0,026 | 0,023 | 0,036 | 0,032 | 0,030 | 0,021 | 0,036 | 0,044 | 0,046 | 0,008 | 0,014 | - |

A phylogenetic tree was constructed using the neighbor-joining method based on Nei’s genetic distance. The tree confirmed the presence of seven groups and separate individual samples belonging to different subspecies and geographical populations (Figure 3). The samples – U1808 Turk-Ugam-u-cau, U1802 Turk-Ugam-u-sos, and U1803 Turk-Ugam-u-cau – formed the clade with largest genetic distances to other samples (Figure S2). Most samples (87) belonged to group III. Groups I and II were represented by subclades.

**Figure 3.**
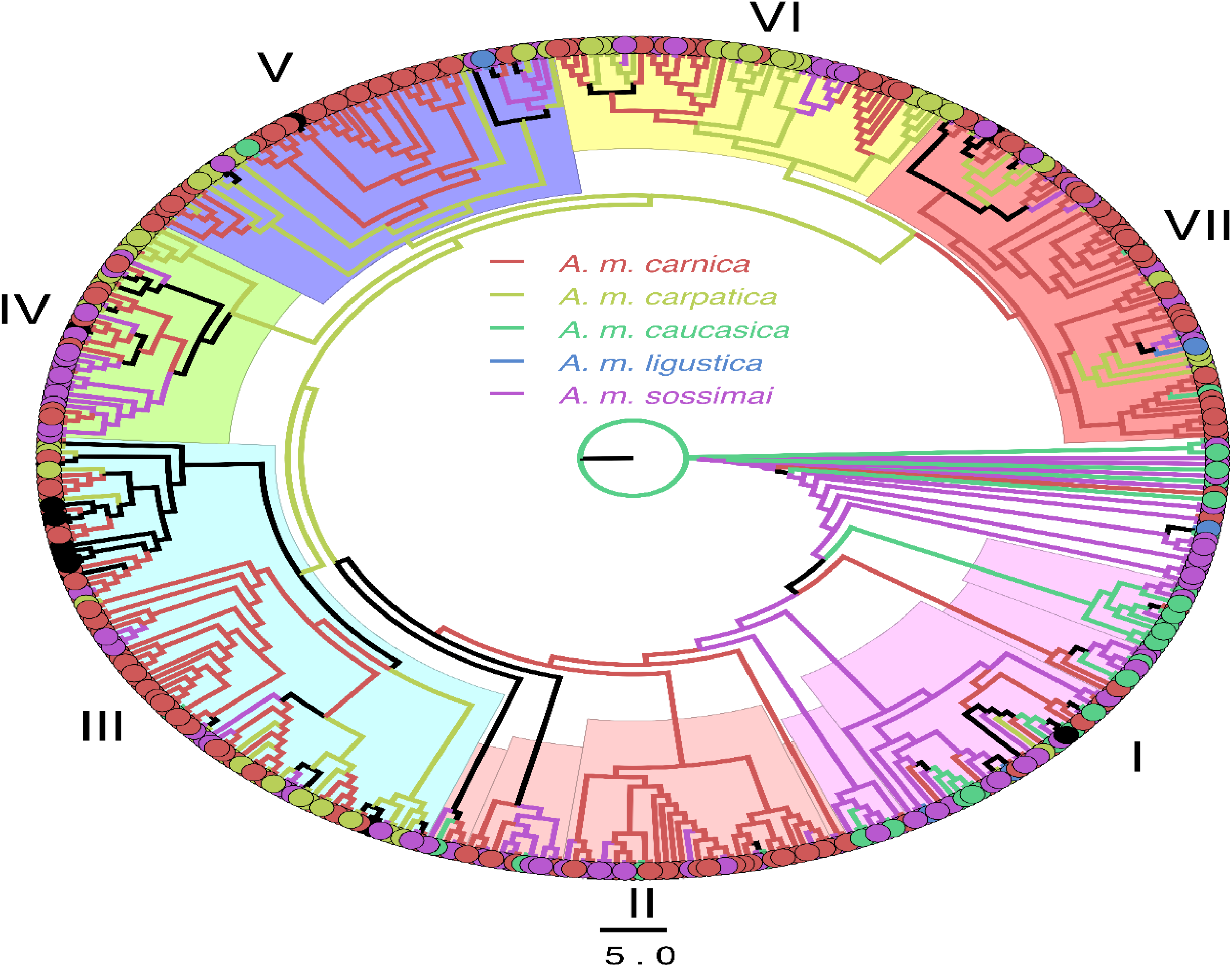
The phylogenetic tree representing the genetic relationship among *Apis mellifera* L. samples that was determined based on Nei’s genetic distance.

### 3.5 Cluster analysis

The diversity of populations showed a high level of heterogeneity according to Bayesian cluster analysis. The cluster analysis confirmed the presence of two clusters (max Δ*K*) (Figure 4). The first cluster comprised five populations, namely the C-group, alm-K-k, car-Zham-A, sos-alm, and sos-alm-Sh-p. The last two populations were represented by *А.m. sossimai* and were located close to one another. The first cluster also included the control group and car-Zham-A *A.m. carnica*, as well as the alm-K-k *A.m.carpatica* population. The second cluster-was represented by nine populations: Alm-Kogaly (*A.m. carnica* and *A.m. ligustica*), Karakonyz-aboriginal-cau-sos, Turk-Tyulbay, Turk-Ugam(u), sos, Zhamb-Karakonyz, Zhet-Alakol (*А.m. caucasica* and *А.m. sossimai*), Zhet-Balambay, and Zhet-Ushbulak-sos. Additionally, ten populations were identified as a mix of the two clusters. The subspecies could not be differentiated based on the clustering because the same subtypes showed different genetic patterns. However, the geographical locations of the populations had a significant influence on the formation of the clusters (Figure 1 and Table 1). The exceptions were the populations from Zhet-Alakol; *А.m. caucasica* and *А.m. sossimai* were assigned to the second cluster while *A.m. carnica* formed a mix of two clusters. Considering that *K* = 5 (Δ*K* = 0.336), only four clusters and eight populations with a single dominant pattern were identified. Nevertheless, *K* = 5 revealed new genetic patterns for the alm-K-k, car-Shym-T, and sos-alm populations. The *A.m. carpatica* specimens of alm-K-k and *А.m. sossimai* of sos-alm-Sh-p differed from the first cluster *A.m. carnica* of *K* = 2. The same trend was observed for *А.m. sossimai* of sos-alm-K. Additionally, one single cluster was represented by the *A.m. carnica* individuals from the car-Shym-T population. The *A.m. carnica* of the car-Zham-A population and *А.m. sossimai* of sos-alm belonged to the same cluster as the samples from the C-group. However, the queens of car-Zham-A and sos-alm were originated from different subspecies and countries, i.e., Germany and Ukraine, respectively. Moreover, the *A.m.carpatica* specimens of alm-K-k and *А.m. sossimai* from the sos-alm-K and sos-alm-Sh-p populations belonged to the same cluster despite having various subspecies and geographical distances of at least 150 km between them. The bee families of these populations were formed by 47 pure lines of *A.m. carpatica* for alm-K-k and pure lines of *А.m. sossimai* for sos-alm. These bee queens originated in Ukraine.

**Figure 4.**
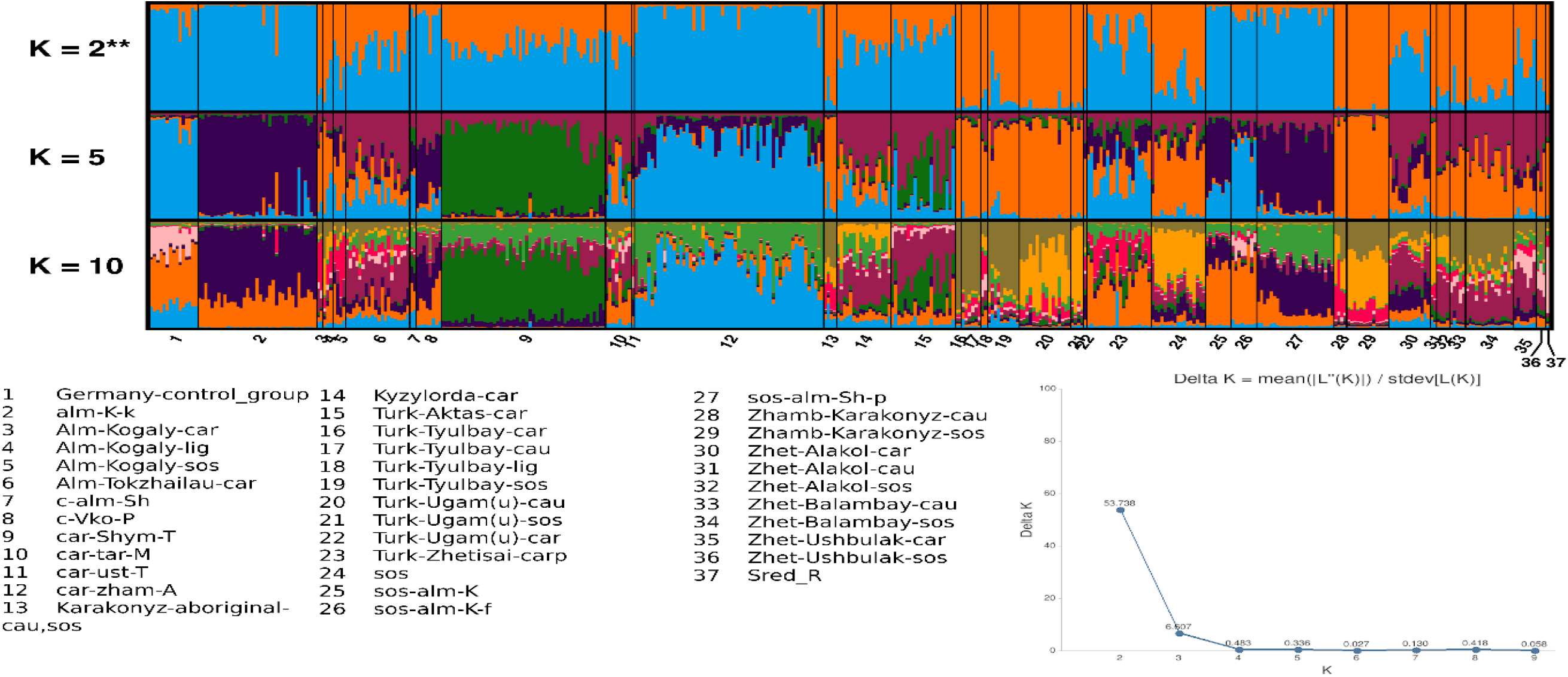
Population structure and genetic differentiation of *Apis mellifera* L. based on microsatellite markers.

### 4. Discussion

The Central Russian bee (*Apis mellifera mellifera*) has been present throughout Kazakhstanian apiaries since its import into Eastern Kazakhstan at the end of the 18^th^ century [6]. *A.m. mellifera* bees were imported to Kazakhstan from Kiev as economically prospective sub-species possessing the ability to breed easily, produce satisfactory amounts of honey, and adapt to the local natural and climatic conditions. Although a disadvantage of this species was their aggressiveness. At present, genetic traces of *A.m. mellifera* can be found in apiaries with other subspecies. The environmental conditions for beekeeping have changed dramatically, partly due to intensive agriculture and the development of animal husbandry, as well as meteorological factors. The natural diversity of plants in some beekeeping zones were replaced by crops, forests became arable lands, and initially humid climates gradually turned to droughts. Several beekeepers thus decided to introduce new bee subspecies and breeds to intensify bee-keeping in Kazakhstan.

The first *Apis mellifera caucasica* populations were imported to Kazakhstan in 1912. The peacefulness and high productivity of the *A.m. caucasica* led to the intensive production of queens at an experimental station (Almaty) in 1930 and in Eastern Kazakhstan in 1950 [6]. At the same time, the imports of *Apis mellifera ligustica* and *Apis mellifera carnica* were registered in several apiaries in Eastern and Southern Kazakhstan. To evaluate the most promising bee subspecies for breeding under the environmental conditions of Kazakhstan, the Kazakh experimental beekeeping station performed bee testing for 11 years starting in 1964 [6]. The results of testing confirmed *A.m. carpatica, А.m. sossimai*, and *А.m. mellifera* as suitable sub-species for breeding.

Since extensive imports of different subspecies began, unregulated bee breeding of the local *A.m. mellifera* with later subspecies laid the foundation to the mixed genetic nature of honeybee populations in Kazakhstan. The apiaries of pure *A.m. mellifera* located in the mountains were also affected by random cross-breeding leading to a decrease in bee productivity and their susceptibility to nosematosis along with other diseases.

The admixture of local honeybee populations was confirmed in this work since the differentiation of these populations through clustering analysis (*K* = 2) (Figure 4) could only be made according to their geographical location. A considerable degree of genetic admixture among subspecies was identified among every studied population. The subspecies were not separated from each other. The statistically confirmed first cluster represented the populations related to the control group, including alm-K-k (*A.m.carpatica*), car-Zham-A (*A.m. carnica*), sos-alm (*А.m. sossimai*), and sos-alm-Sh-p (*А.m. sossimai*). These three subspecies belonged to the same genetic cluster according to *K* = 2. The car-Zham-A and sos-alm-K-f populations maintained the same genetic pattern as the control group in *K* = 5 while sos-alm-K (Kargaly) and sos-alm-Sh-p (Shelek) formed a new cluster with alm-K-k (Kokpek). The geographical distances between Kargaly and Shelek and Kokpek, Shelek, and Kokpek were at least 150 km and 35 km, respectively. The maximum recorded mating distance for the honeybees was 15 km [25,26]. The genetic relationship could possibly be explained by bee material exchange between beekeepers without registration or mention in notes of apiaries. Nevertheless, it should be highlighted that this pattern was formed by two different subspecies, namely *A.m. carpatica* and *А.m. sossimai*. In addition, the *A.m. carpatica* of the c-Vko-P population located in Ulan (Eastern Kazakhstan) showed a similar pattern as the later populations but their distance to the nearest population of sos-alm-Sh-p was around 500 km. The unique cluster (*K* = 5) was presented by *A.m. carnica* from the car-Shym-T population located in Shymket. The apiary represented by the car-Shym-T population was established in 2018 by a bee queen belonging to the Troisek line from Germany. Breeding within this apiary continues to be supported by its own queens without the addition of new lines or breeds. The most genetically structured populations (*K* = 2, 5) consisted of Zhamb-Karakonyz (*А.m. sossimai, А.m. caucasica*), Turk-Ugam(u) (*А.m. sossimai, А.m. caucasica, A.m. carnica*), and Turk-Tyulbay (*А.m. sossimai, А.m. caucasica, A.m. carnica, A.m. ligustica*). The latter population included four subspecies without any differentiation by type. According to the cluster analysis, most populations were formed by uncontrolled breeding within populations. In addition, excess outbreeding was confirmed for the car-Shym-T, car-zham-A, Zhet-Alakol, and Zhet-Ushbulak populations which comprised more heterozygotes than expected. The outbreeding which may have happened with the nearby apiaries was not investigated in this research. The other 12 populations showed a high level of admixture but remained the evolution line C. The samples belonged to the C line with the exception of two samples of *А.m. sossimai* and four samples of *A.m. carnica* from the Zhet-Ushbulak population; these were represented by the M4 haplotype like *А.m. mellifera* specimen collected from the Belukha mountain at an altitude 1800 m. Cross-breeding with *А.m. mellifera* could have taken place because the abandoned apiaries populated by *А.m. mellifera* located near to well-maintained ones [6]. In addition, according to the phylogenetic analysis, seven groups consisted of admixtures of different subspecies (Figure 3, Figure S2). The first group comprised four subclades without any differentiation according to their geographical locations or subtypes; indeed, samples from 15 populations were included in group I. Group II also included four subclades of different subspecies from 14 populations. These two groups were more diverse than the other five groups. Unlike the other groups, group V included most samples of *A.m. carnica* from a single population from car-zham-A; this group contained four subclades including two samples of *A.m. carnica* from the control group. Group VI included subclades consisting of samples from the car-Shym-T, alm-K-k, car-zham-A, and c-Vko-P populations. Most samples consisted of *A.m. carnica* and *A.m. carpatica*. None of the formed groups included any precise subspecies or populations and instead, all groups represented an admixture of populations. The populations could not be compared with *А.m. mellifera* and an introgression coefficient could not be determined since only one sample from the Belukha mountain was available. Moreover, the genetic purity of the Central Russian bee could not be confirmed. At present, no reference populations of any subspecies or breeds are formed in Kazakhstan; most apiaries are private, and therefore, it is almost impossible to determine the exact origin of admixed bee populations. Additionally, the Tien Shan Mountains (Kazakhstan side), where most of the studied specimens were collected, are a region of endemic honeybee distribution (*Apis mellifera pomonella*) [27], (Figure 1). It could be concluded that some samples were the result of cross-breeding with *A.m. pomonella* since beekeepers commonly do not maintain pure breeding as was shown for the Zhet-Ushbulak population which included both the C and M lineages.

The absence of government confirmed standards to determine the purity of bee sub-species or breeds; uncontrolled breeding on experimental stations and apiaries; the extensive import of queens of different subspecies and breeds without registration in publicly available resources or in government databases; abandoned apiaries without support from the government; the absence of genetic profiling for queens and records on breeding are the main problems harming the beekeeping in Kazakhstan. To the best of our knowledge, this was the first research focused on investigating the genetic structure of bee populations distributed in the largest apiaries of the country which are included in the government alliance for maintaining beekeeping. Our results suggest that the promiscuous genetic nature of honeybee populations in Kazakhstan is the result of cross-breeding that has been occurring for around 50 years.

## 5. Conclusions

According to our findings, honeybee populations in Kazakhstan are admixtures of different subspecies and breeds separated only by geographical location according to cluster analysis. The constructed phylogenic tree revealed seven groups without separations by subspecies or geographical locations. The revealed promiscuous genetic nature of honeybee populations in Kazakhstan is the result of cross-breeding which has been taking place for around 50 years.

## Supplementary

Table S1. Raw data of bee sample genotyping using microsatellite markers

Table S2. Nei’s genetic distance among the geographically divided populations

Figure S1. The results of DraI mtDNA COI-COII test

Figure S2. The cladogram representing the genetic relationship among *Apis mellifera* L. samples

