## Supplementary material for "Genetic investigation of honeybee populations in Kazakhstan": Figure S1: Figure S1.pdf

DraI mtDNA COI-COII test

016 018 019 020 243 244 245 001 002 003 004 005 M 242 006 007 008 009 010 011 012 013 014 015

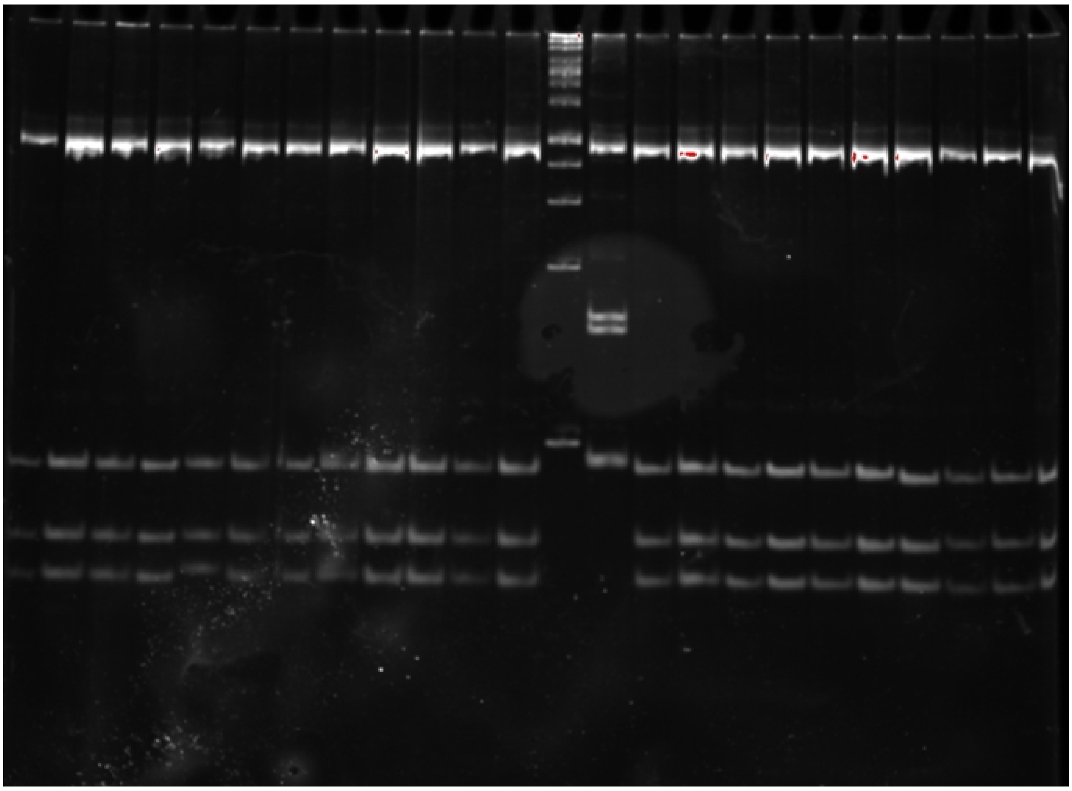

022 024 027 029 030 033 034 035 036 039 K+(243) M K+(242) 040 041 042 043 044 045 046 047 048 049

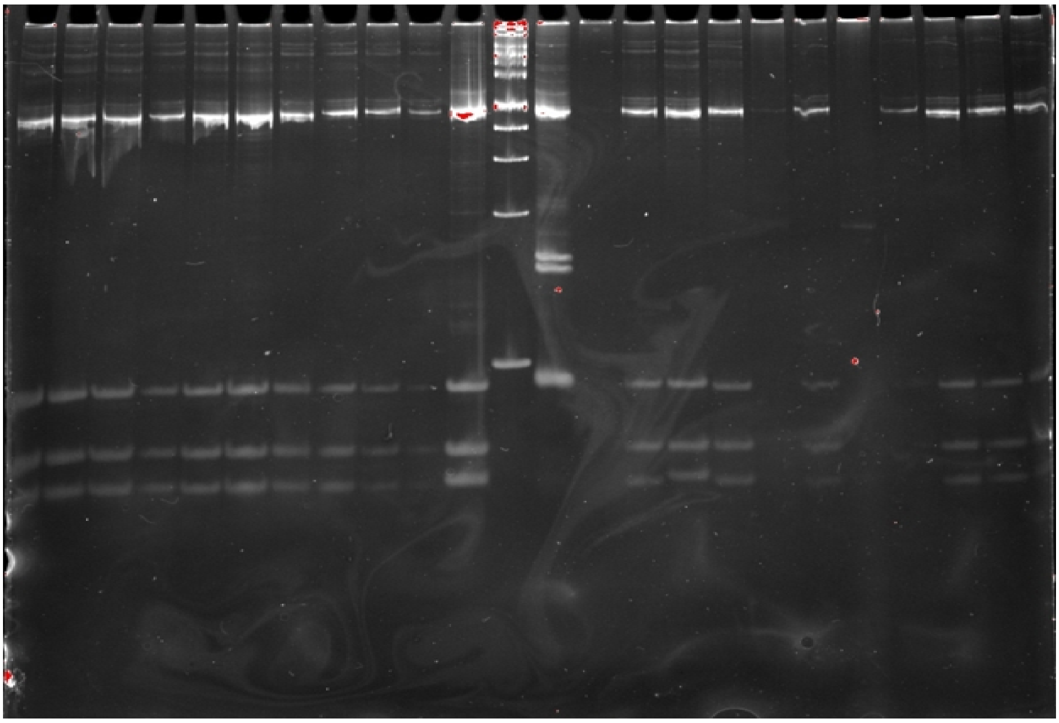

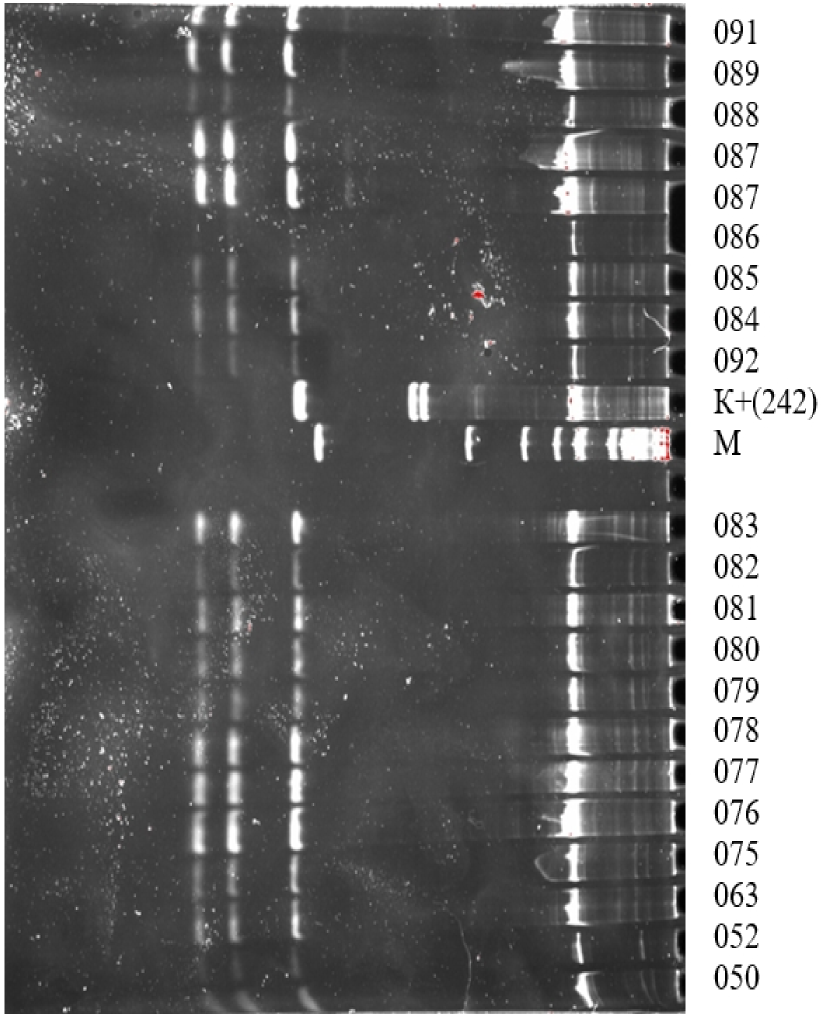
091
089
088
087
087
086
085
084
092
K+(242)
M
083
082
081
080
079
078
077
076
075
063
052
050

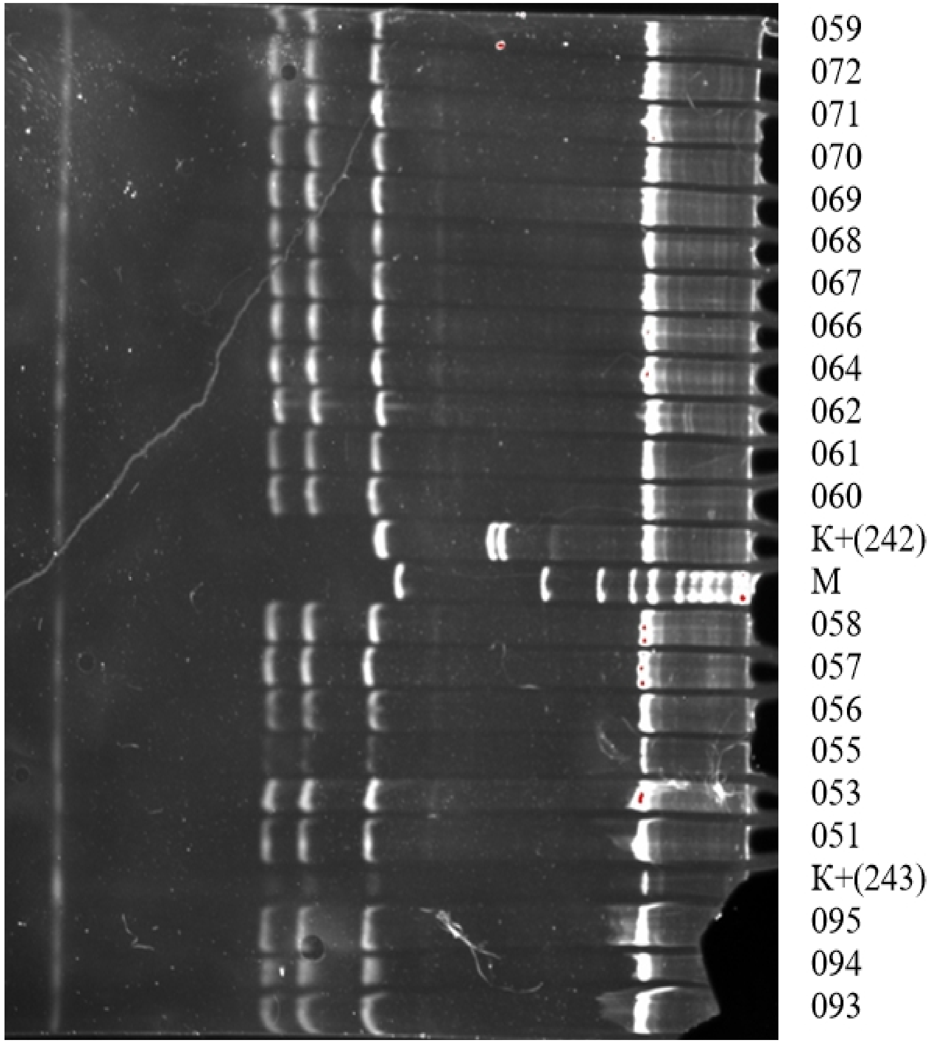
059
072
071
070
069
068
067
066
064
062
061
060
K+(242)
M
058
057
056
055
053
051
K+(243)
095
094
093

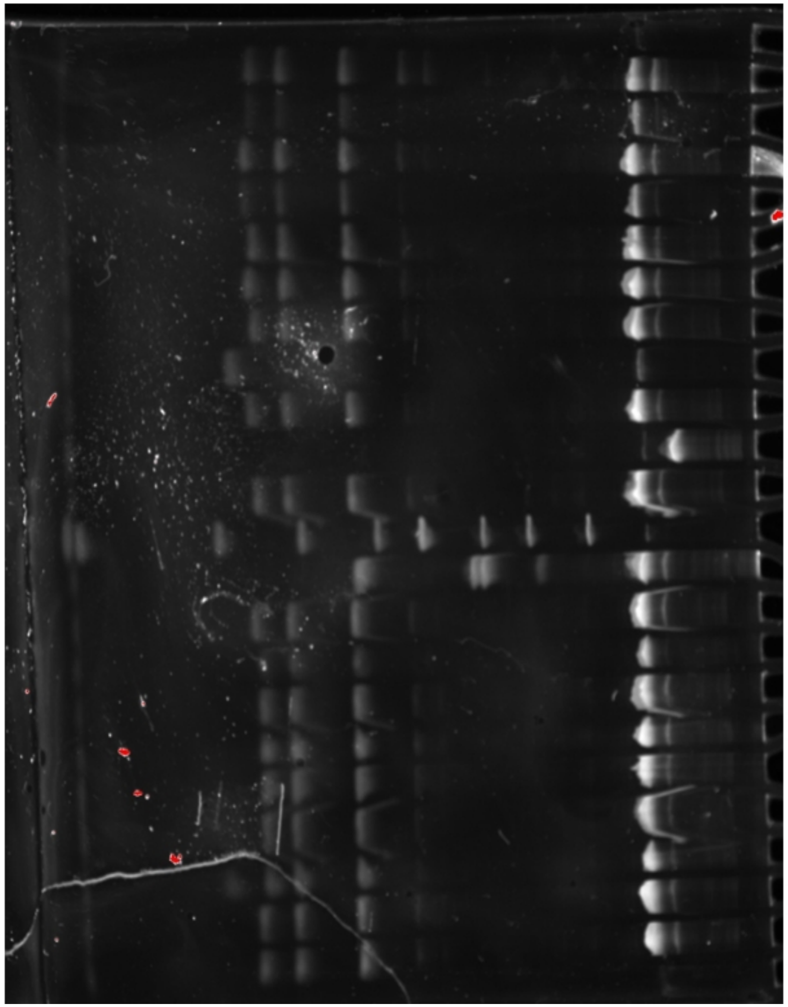
K+
186
074
096
097
098
100
101
179
M
K+ (243)
K+ (242)
102
103
191
189
188
032
037
038
043
045

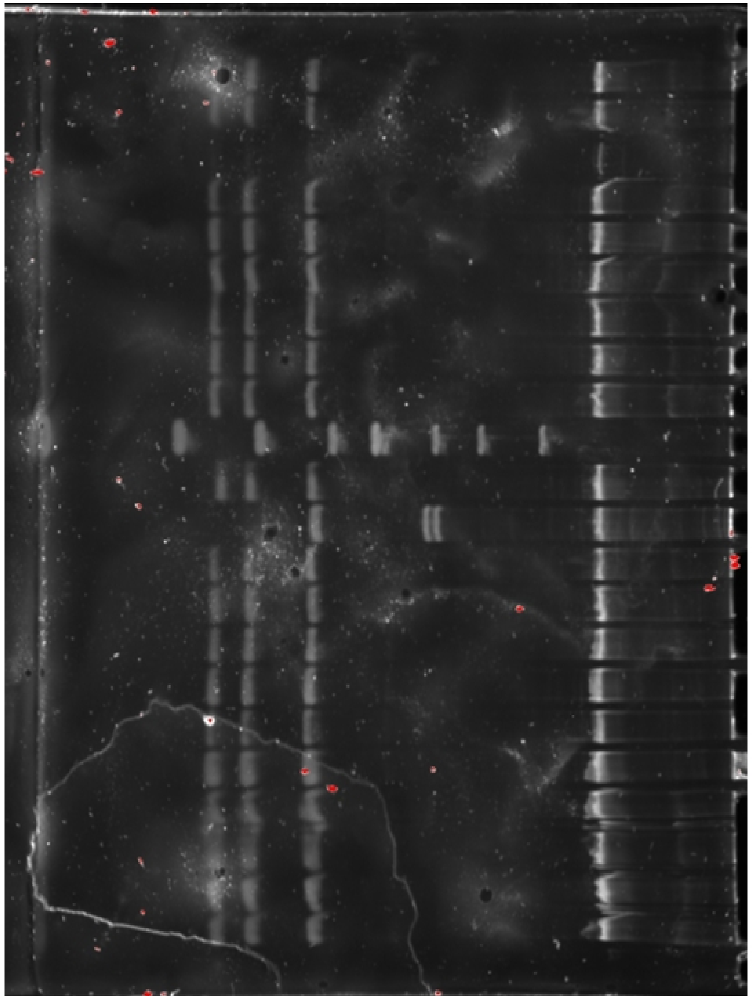
148
139
135
134
129
125
122
115
114
M
K+(243)
K+(242)
113
112
111
110
109
107
106
105
104
065

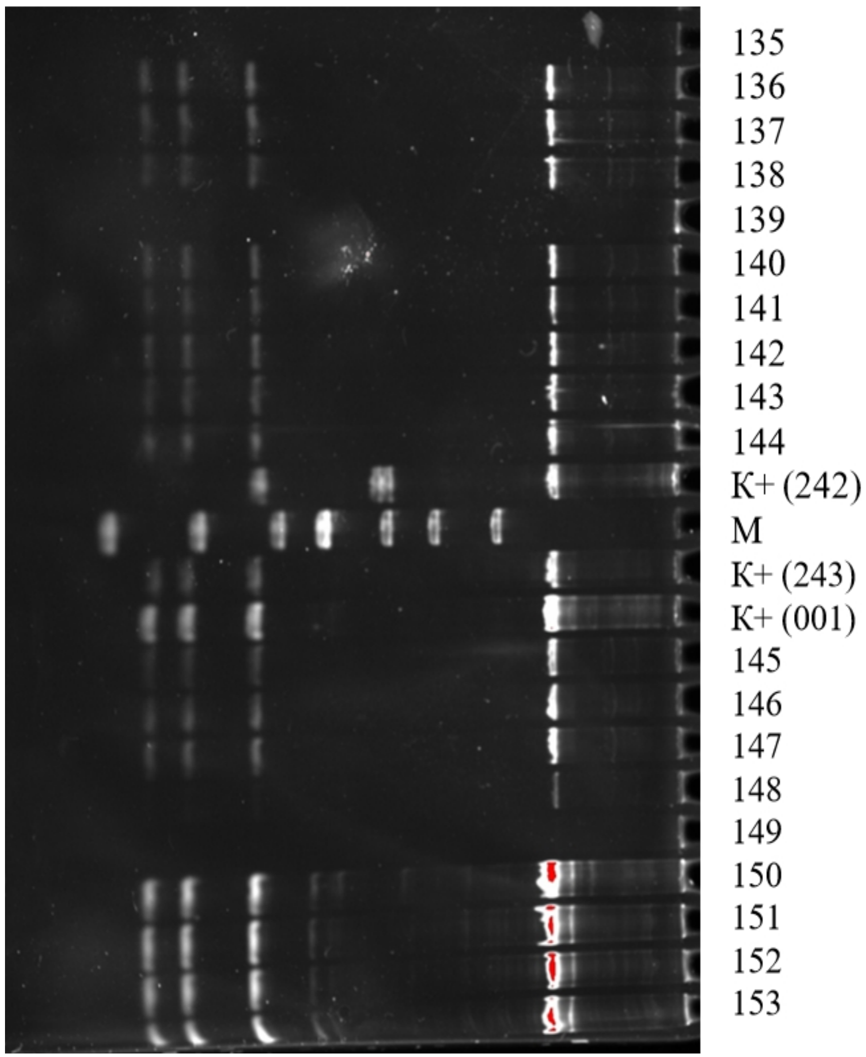
K+ (242)
M
K+ (243)
K+ (001)
145
146
147
148
149
150
151
152
153

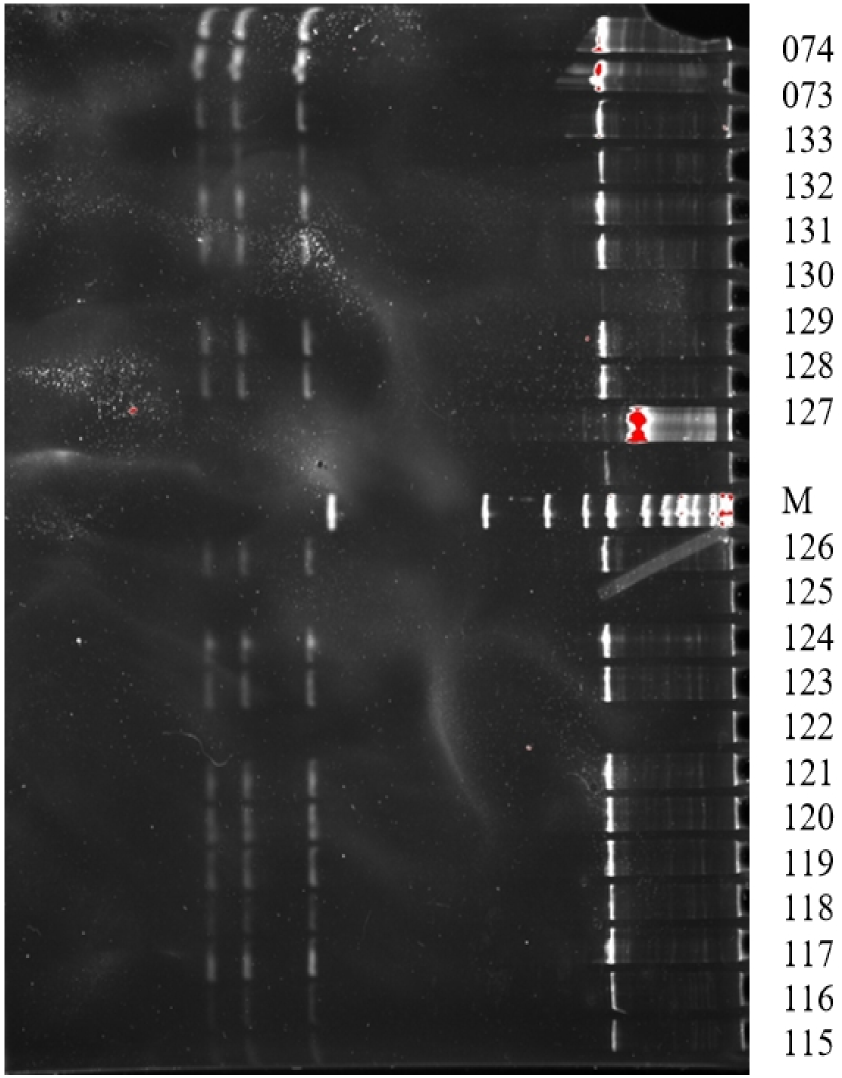
074
073
133
132
131
130
129
128
127
M
126
125
124
123
122
121
120
119
118
117
116
115

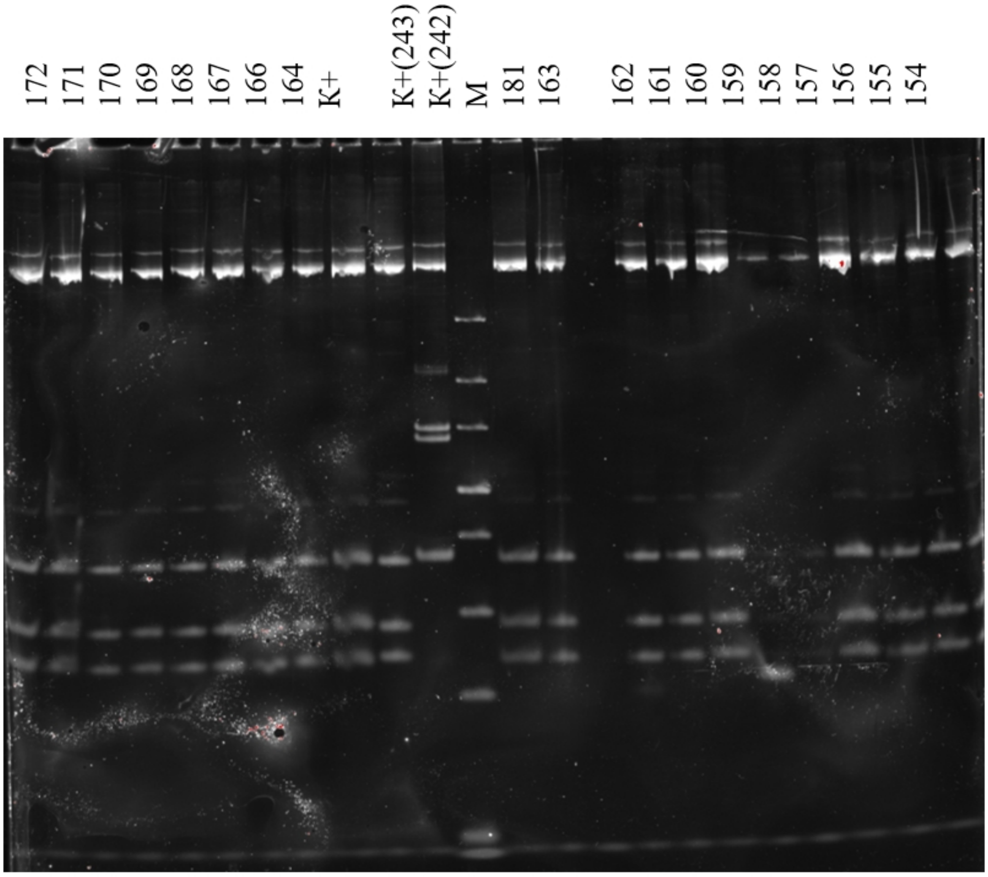
172
171
170
169
168
167
166
164
K+
K+(243)
K+(242)
M
181
163
162
161
160
159
158
157
156
155
154

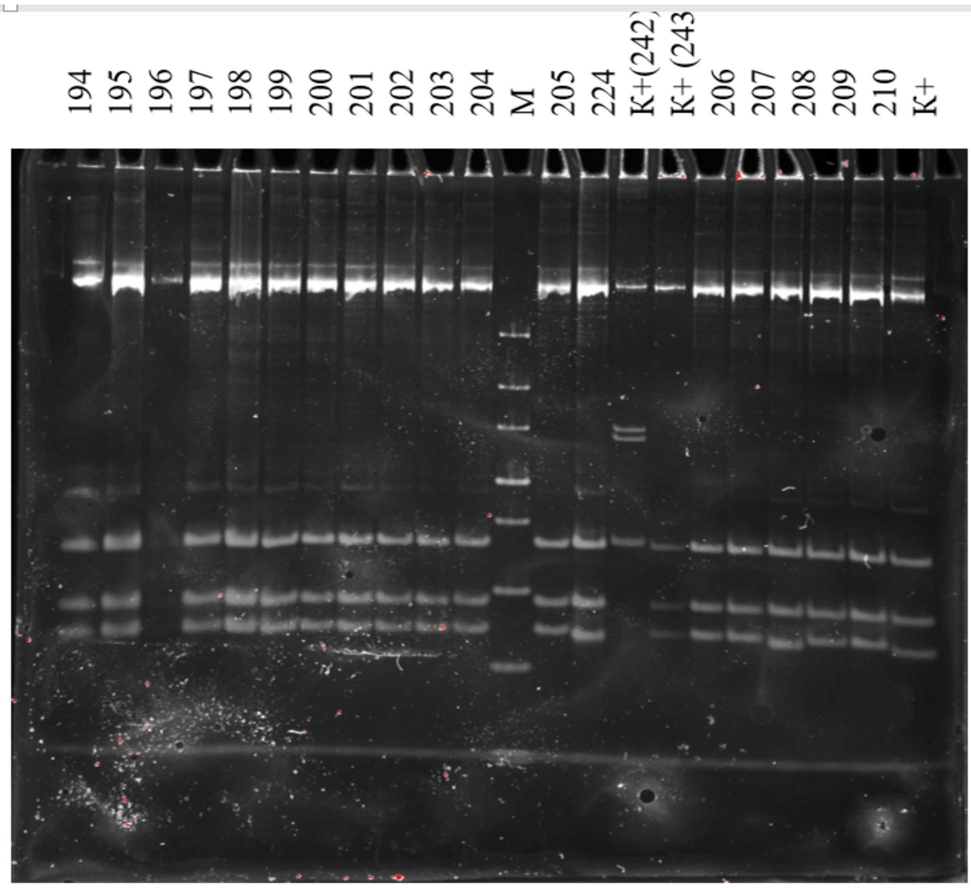
M
205
224
K+(242)
K+ (243
206
207
208
209
210
K+

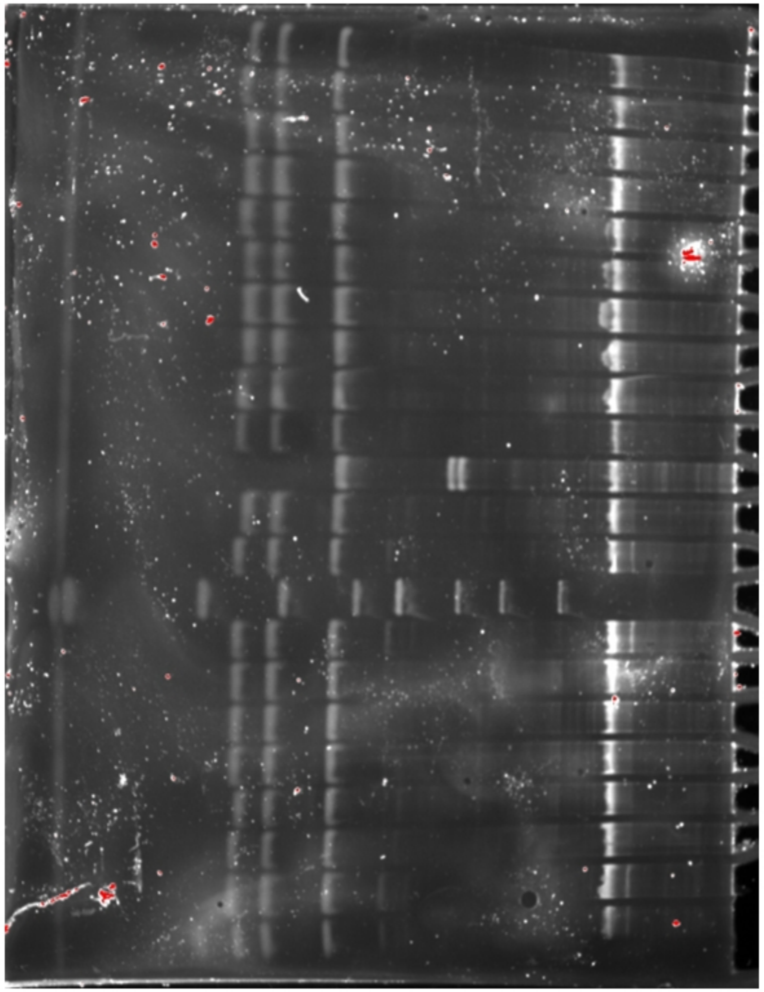
K+(242)
K+(243)
K+(1)
M
236
237
238
240
239
241
174
175

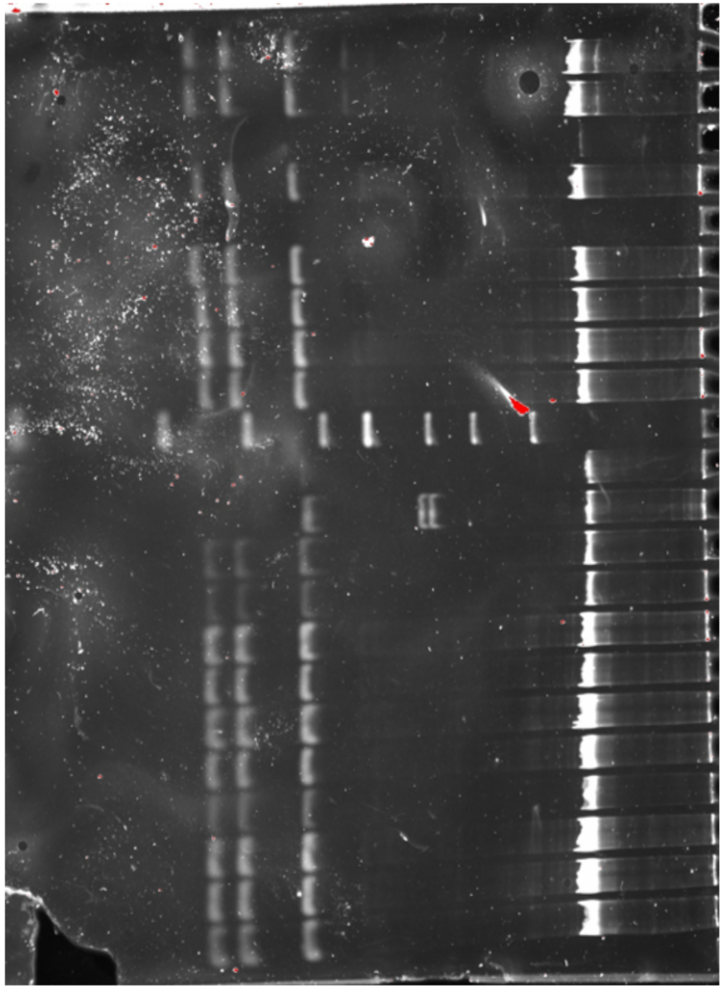
K+
148
225
224
223
222
221
220
M
K+(242)
K+(243)
219
218
217
216
215
214
213
212
211

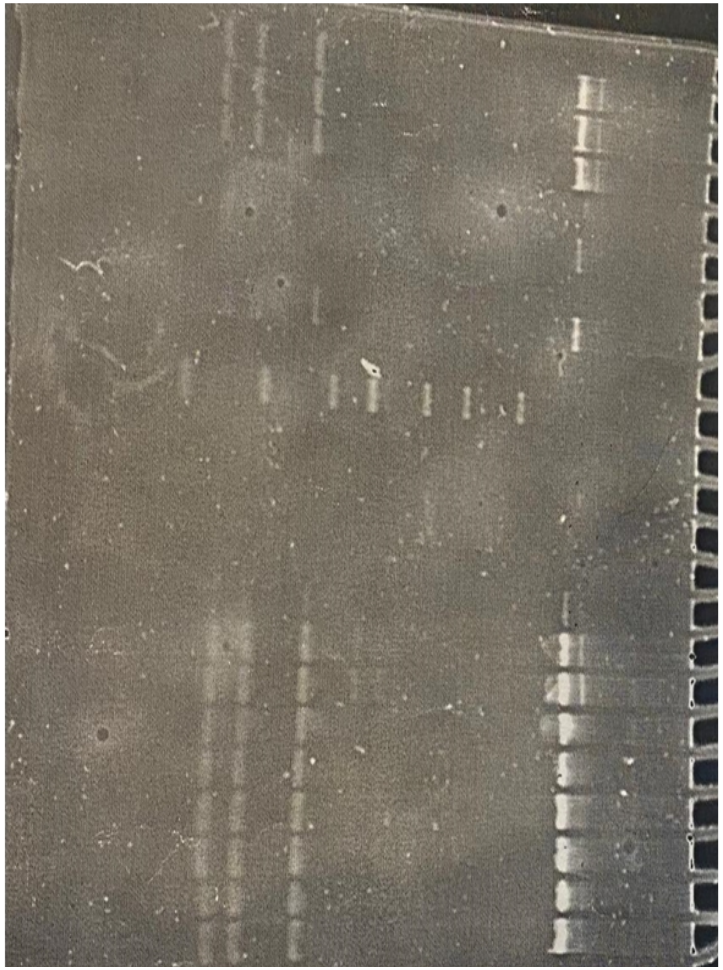
172
171
164
M
090
054
T116
T108
K1516
T115
S1145
S1011

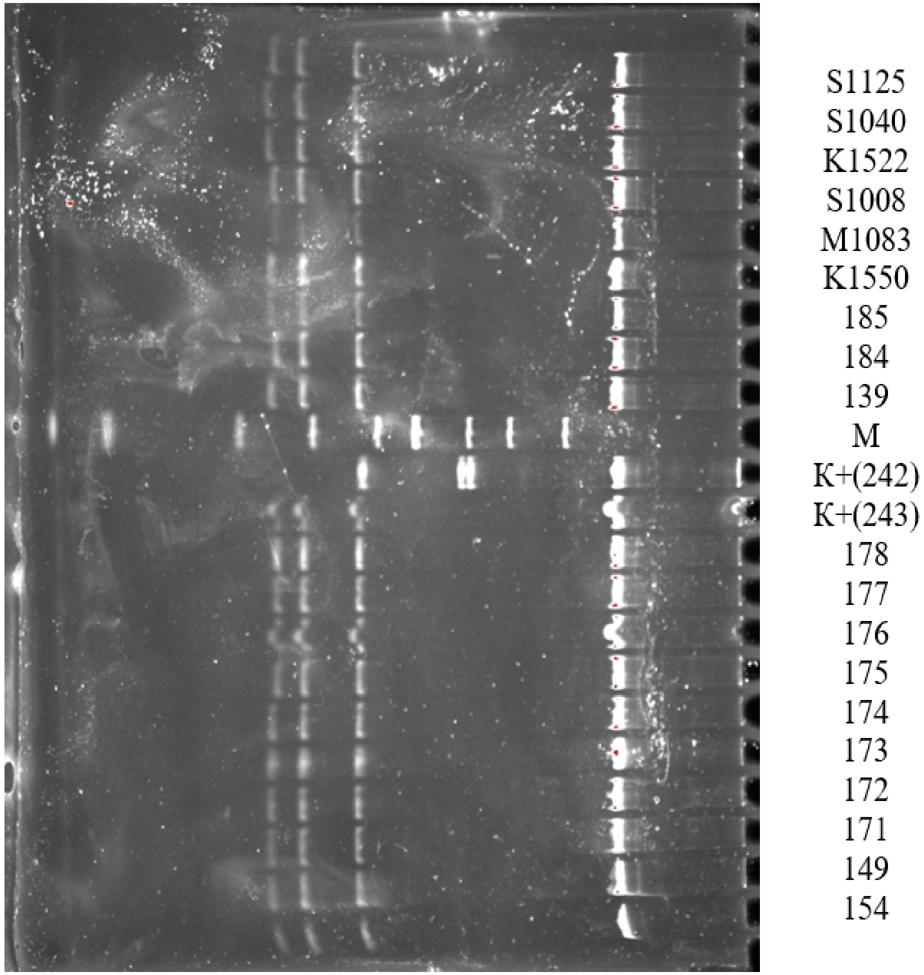
S1125
S1040
K1522
S1008
M1083
K1550
185
184
139
M
K+(242)
K+(243)
178
177
176
175
174
173
172
171
149
154

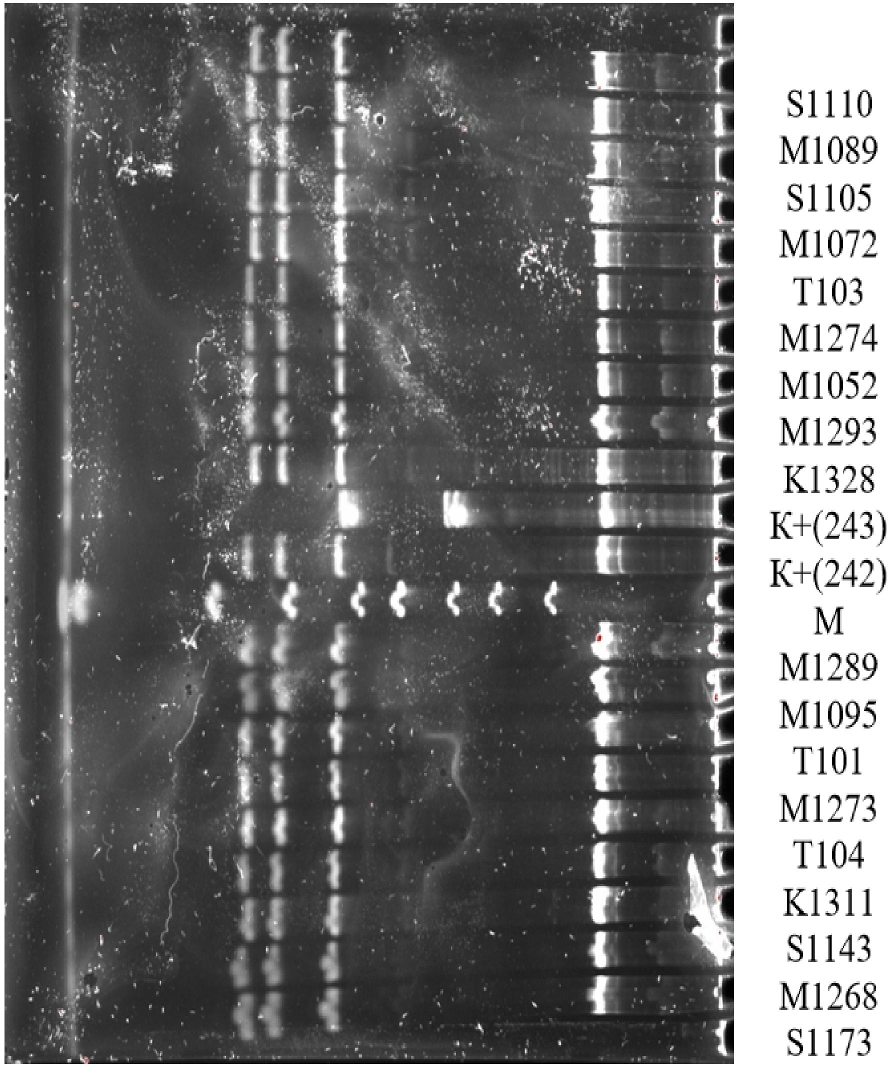
S1110
M1089
S1105
M1072
T103
M1274
M1052
M1293
K1328
K+(243)
K+(242)
M
M1289
M1095
T101
M1273
T104
K1311
S1143
M1268
S1173

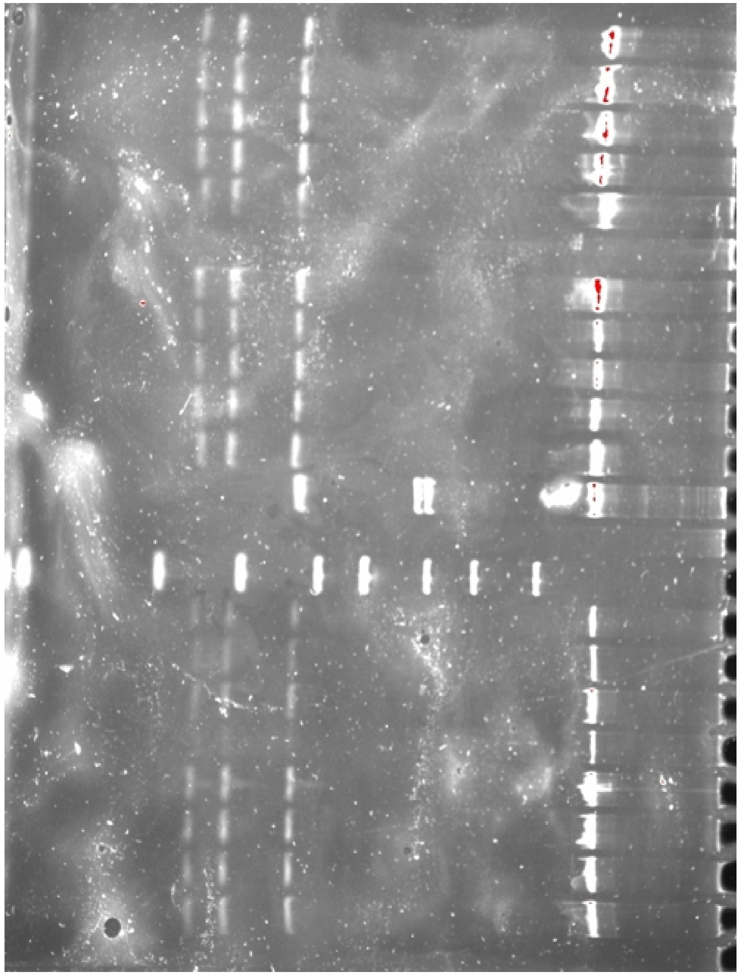
M1064
M1066
S1112
K1543
S1014
M1091
M1087
K1344
T107
T109
K+(243)
K+(242)
M
G5
G11
G14
G22
K2
K4
K5
K10

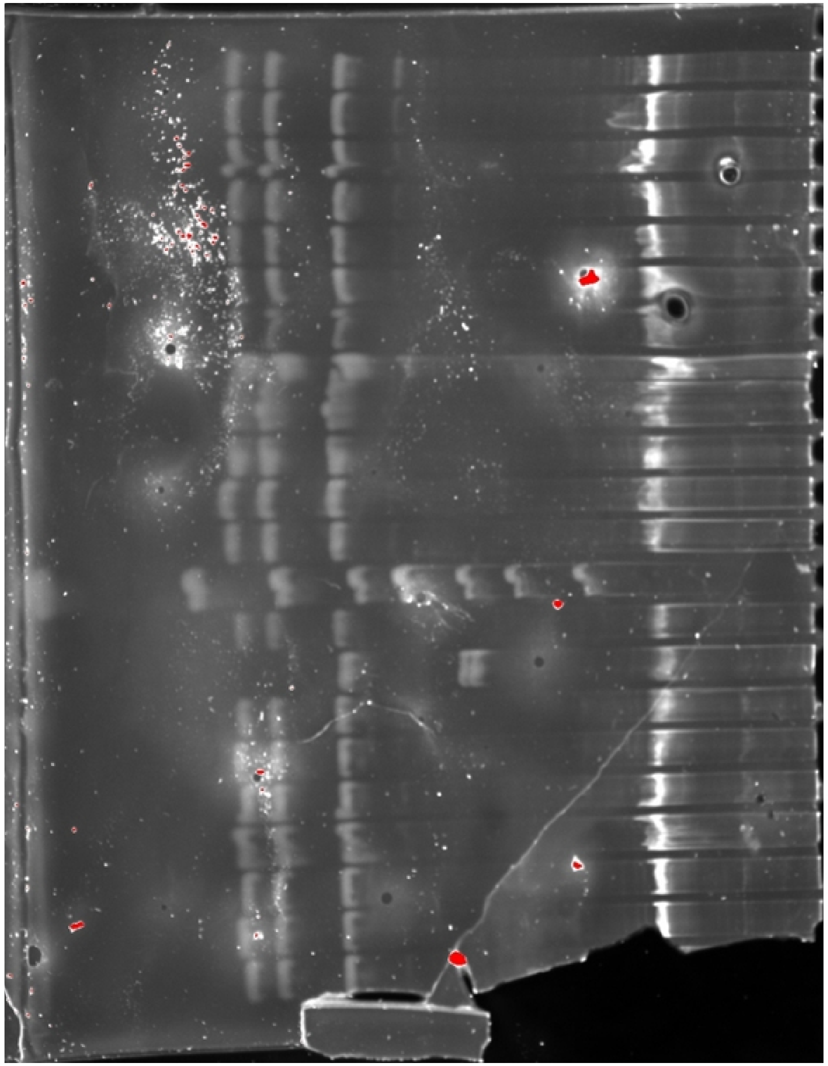
K1324
K10
K1550
M1083
K1535
T117
T111
K1307
S1117
T103
T106
K1304
M
K+(243)
K+(242)
S1011
K1522
S1008
K1546
S1013
M1066
K1350

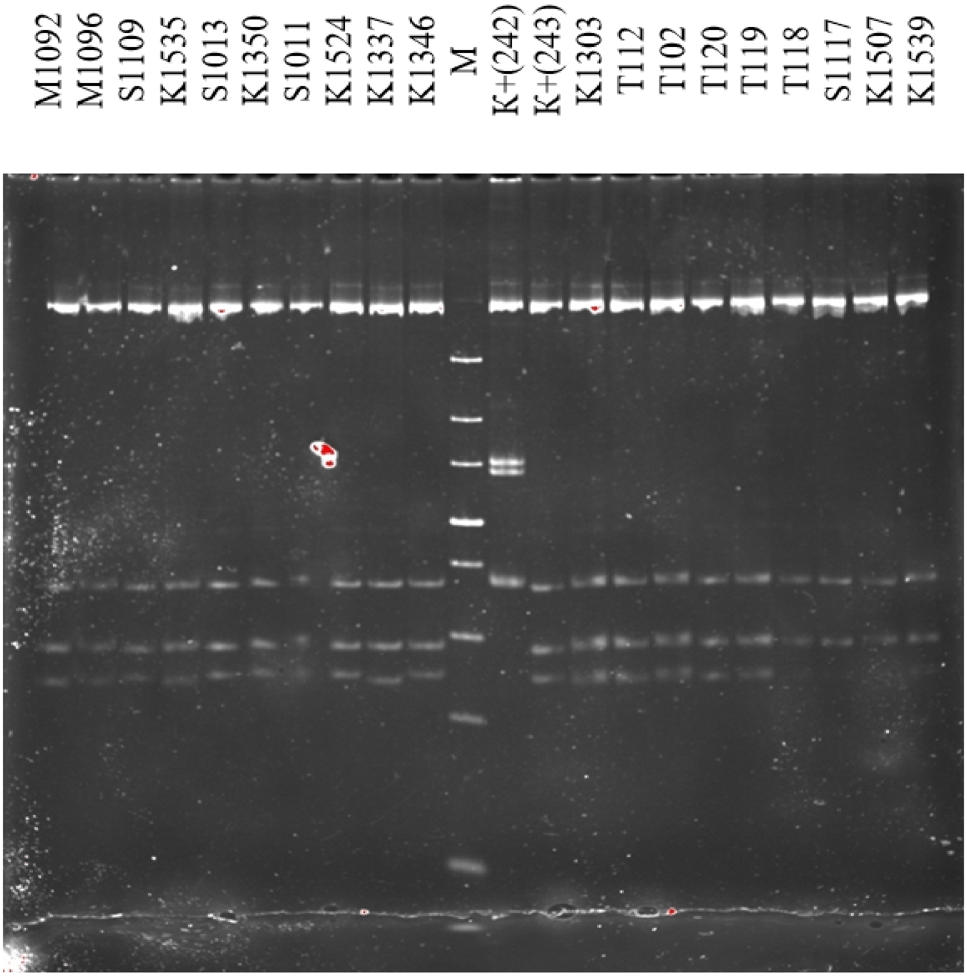
M1092
M1096
S1109
K1535
S1013
K1350
S1011
K1524
K1337
K1346
M
K+(242)
K+(243)
K1303
T112
T102
T120
T119
T118
S1117
K1507
K1539

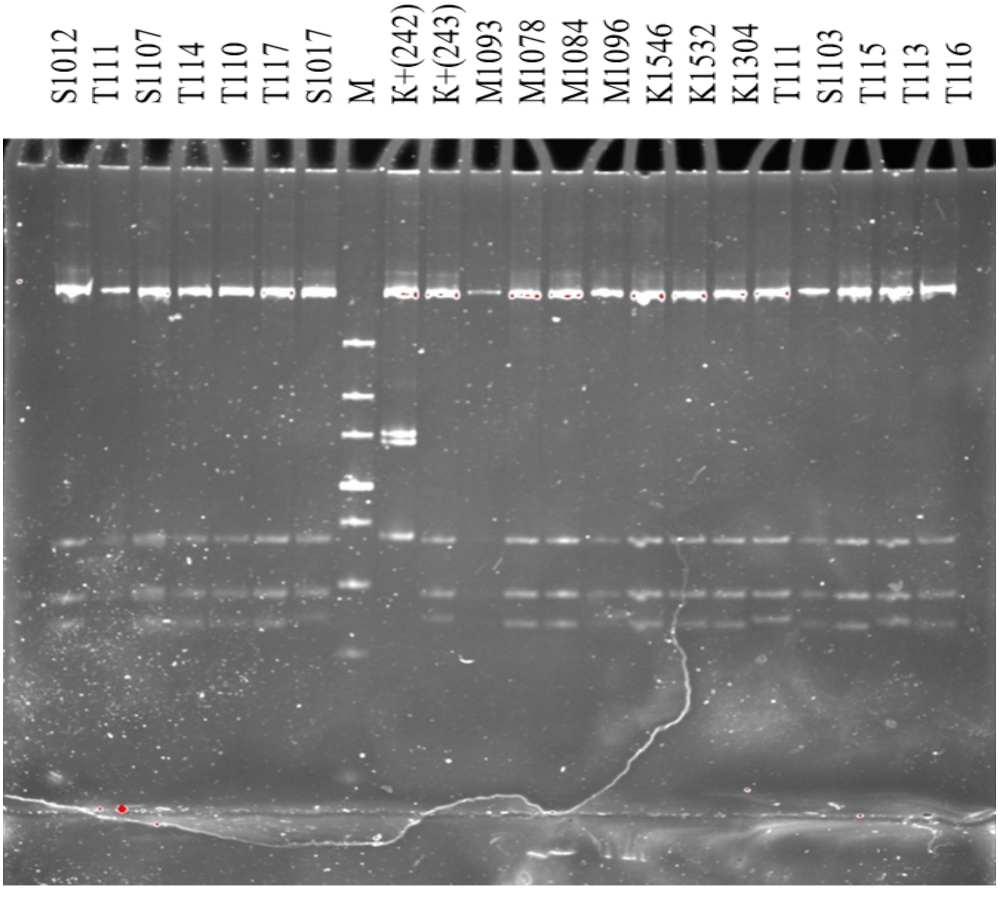
S1012
T111
S1107
T114
T110
T117
S1017
M
K+(242)
K+(243)
M1093
M1078
M1084
M1096
K1546
K1532
K1304
T111
S1103
T115
T113
T116

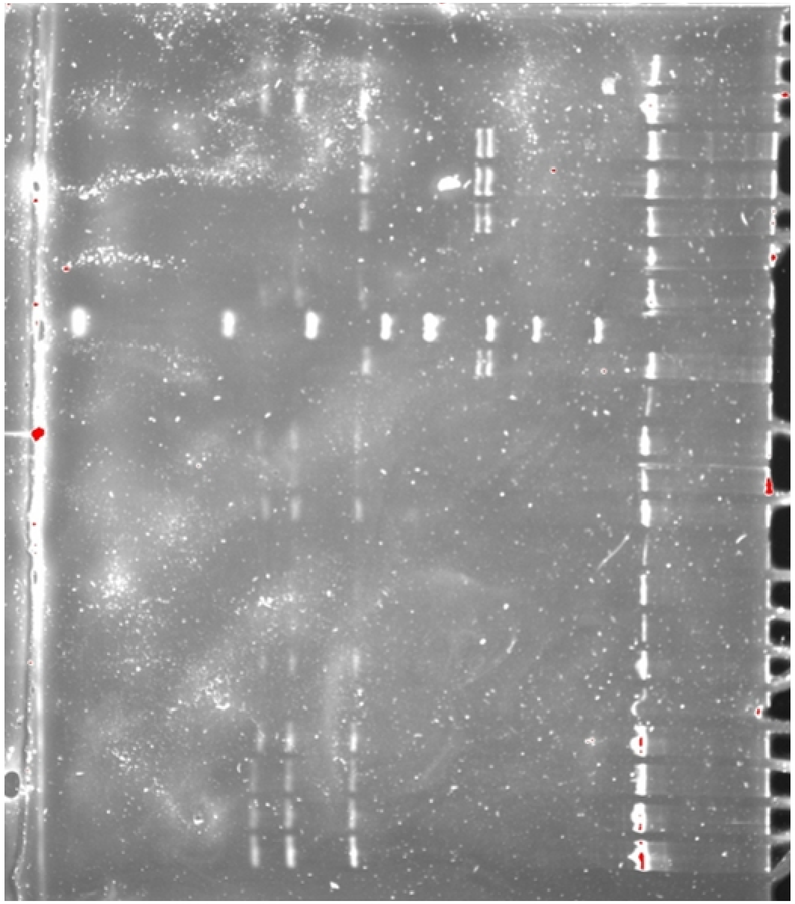
N17
F4
F10
F6
F8
U1802
1807
M
K+(242)
K+(243)
U1808
G33
G25
G17
G18
G16
G15
G6
G4
K16
K12
K3

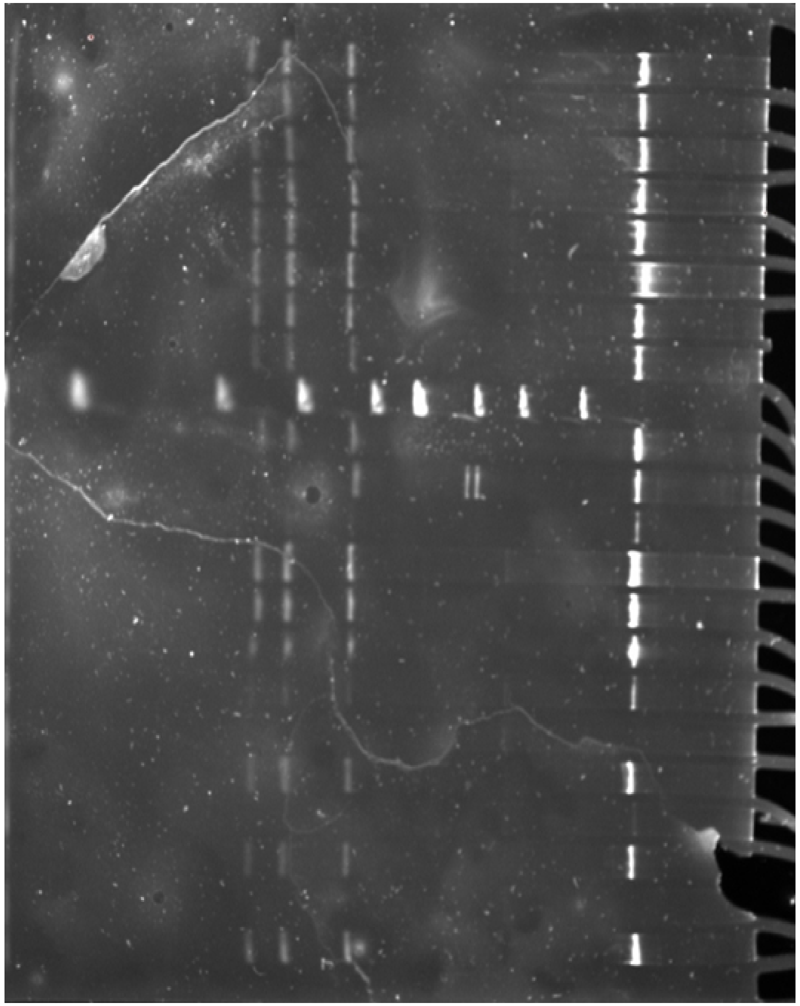
U1805
U1801
U1809
U1812
U1811
U1813
U1806
U1804
M
K+(243)
K+(242)
G36
G44
G12
T106
M1057
S1124
K1312
M1091
K1307
S1010
T105

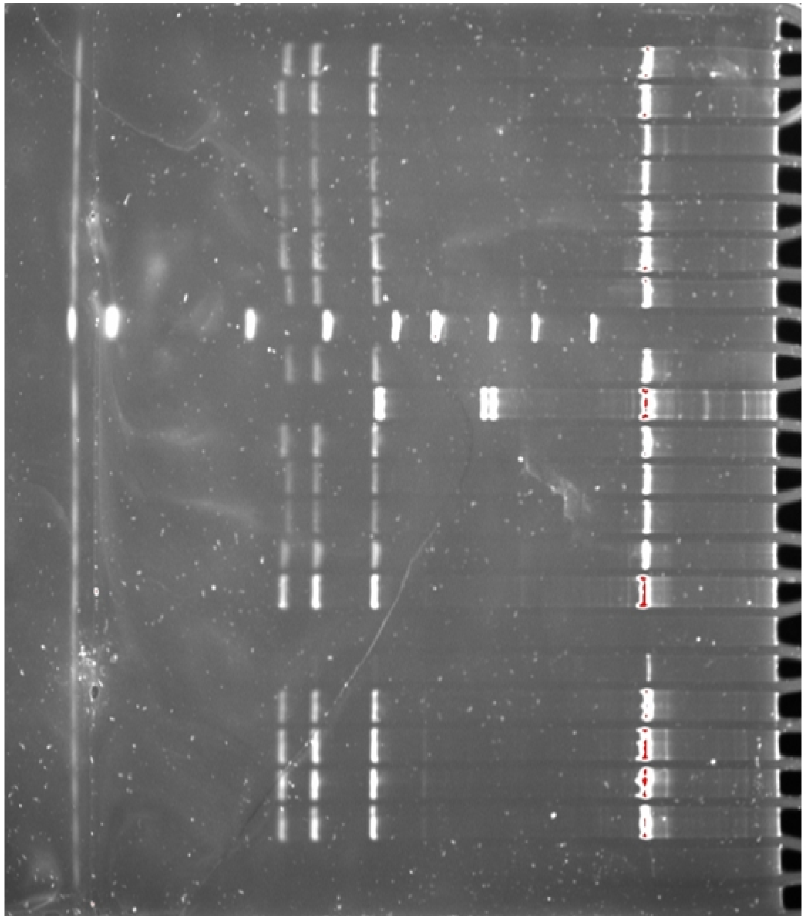
A130
A129
A121
A108
A116
A117
A131
M
K+(243)
K+(242)
A119
G10
G17
A162
A118
SPK21
N55
N127
N19
N57
N97
SPK1

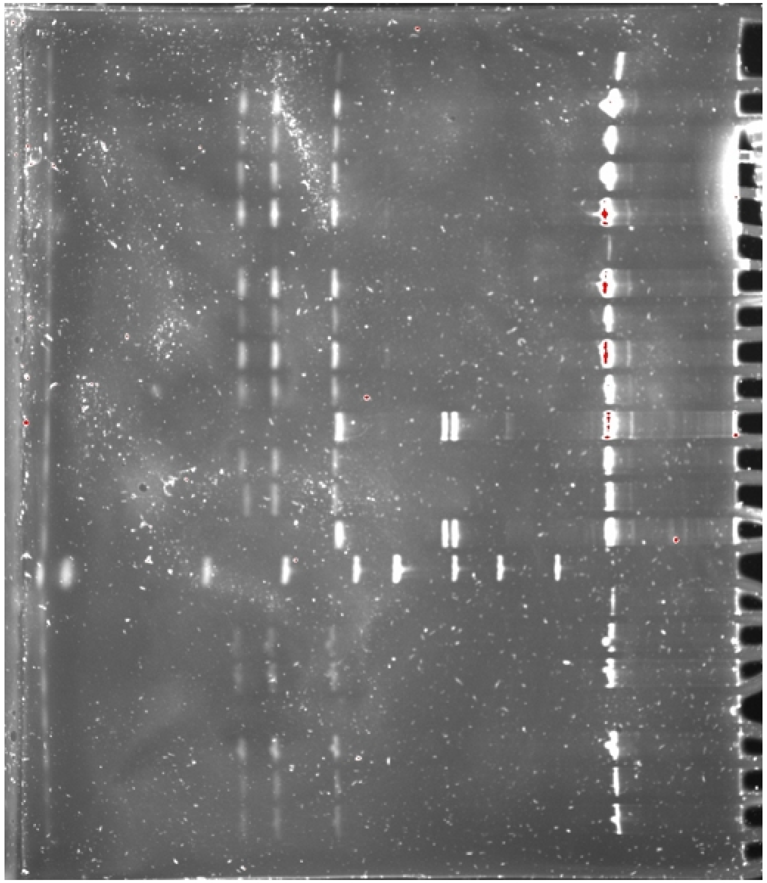
U1808
TN1
TN2
TN3
TN4
TN5
U1814
U1816
U1815
F1
F5
SPK6
K+(243)
K+(242)
M
SPK8
SPK5
SPK3
N12
N17
N4
N15

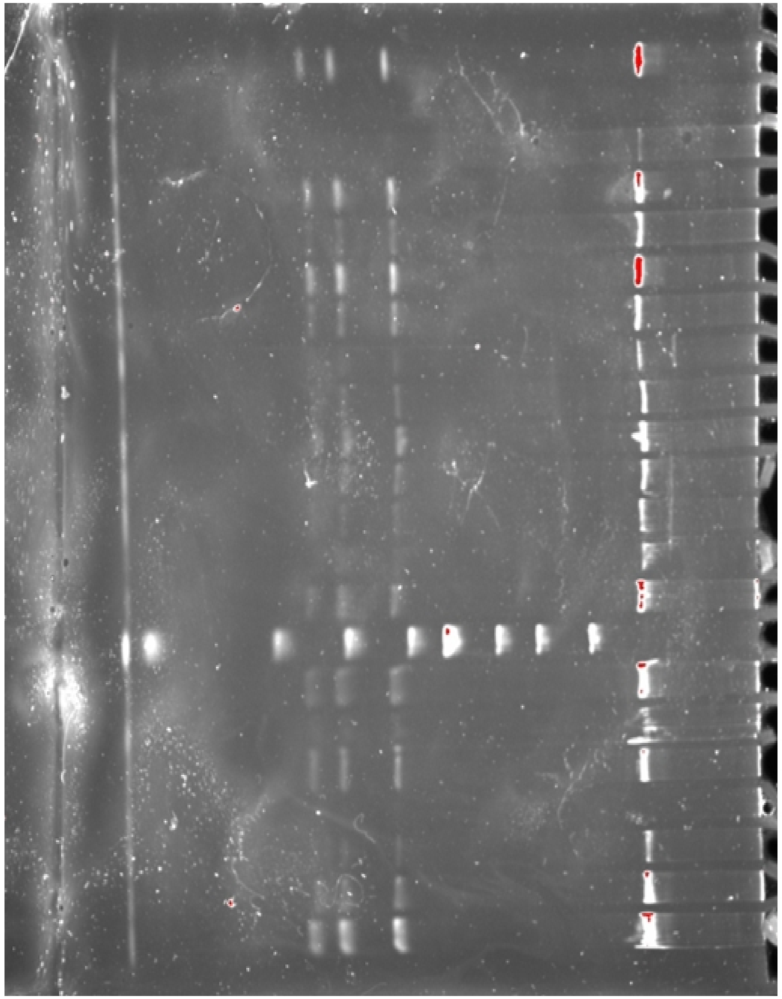
SPK19
SPK16
SPK18
SPK11
SPK10
SPK13
SPK9
SPK12
SPK15
SPK17
K13
K18
K+(243)
M
A167
A122
SPK20
SPK7
SPK8
A138
K14

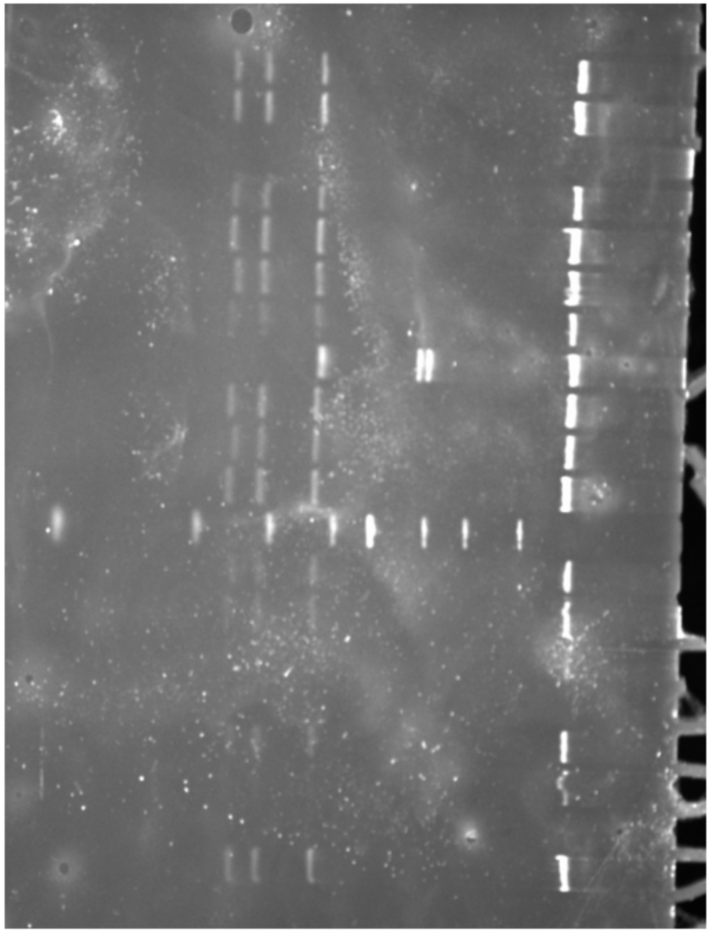
SPK1
SPK2
SPK14
SPK10
G16
SPK7
K+(242)
SPK18
K+(243)
K18
M
A120
A188
SPK21
N55

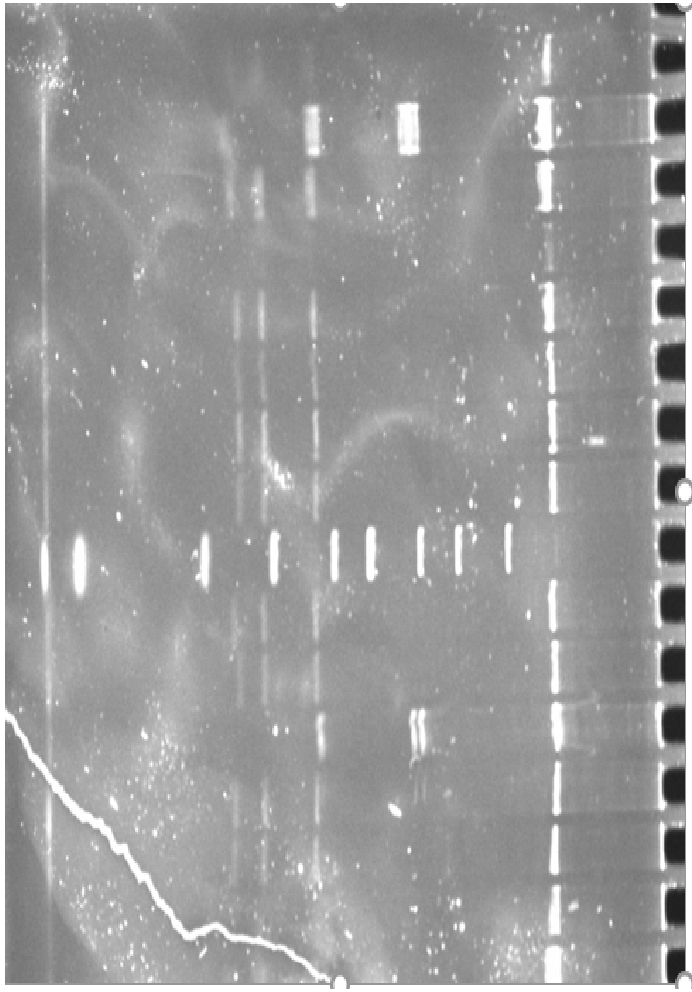
N6
F7
F2
G16
G28
G3
G15
M
S1124
K+(243)
K+(242)
U1807
U1810

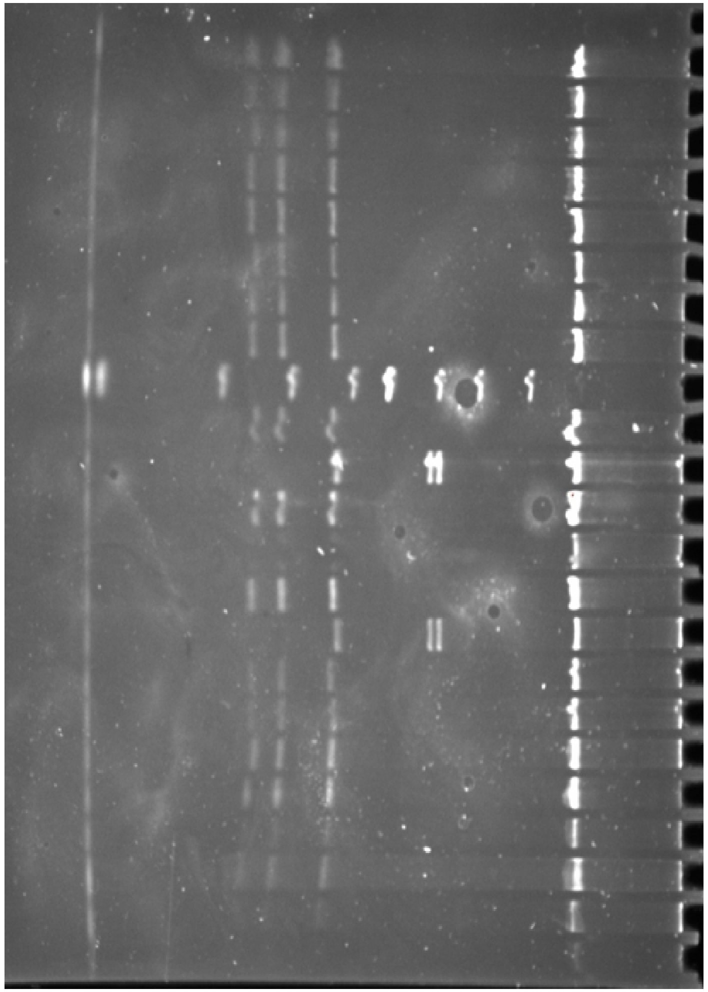
A136
N6
A132
A128
F3
S1010
U1802
M1091
M
K+(243)
K+(242)
A171
SPK17
K13
F9
A122
TN5
A173
SPK16
SPK15
N127
A188

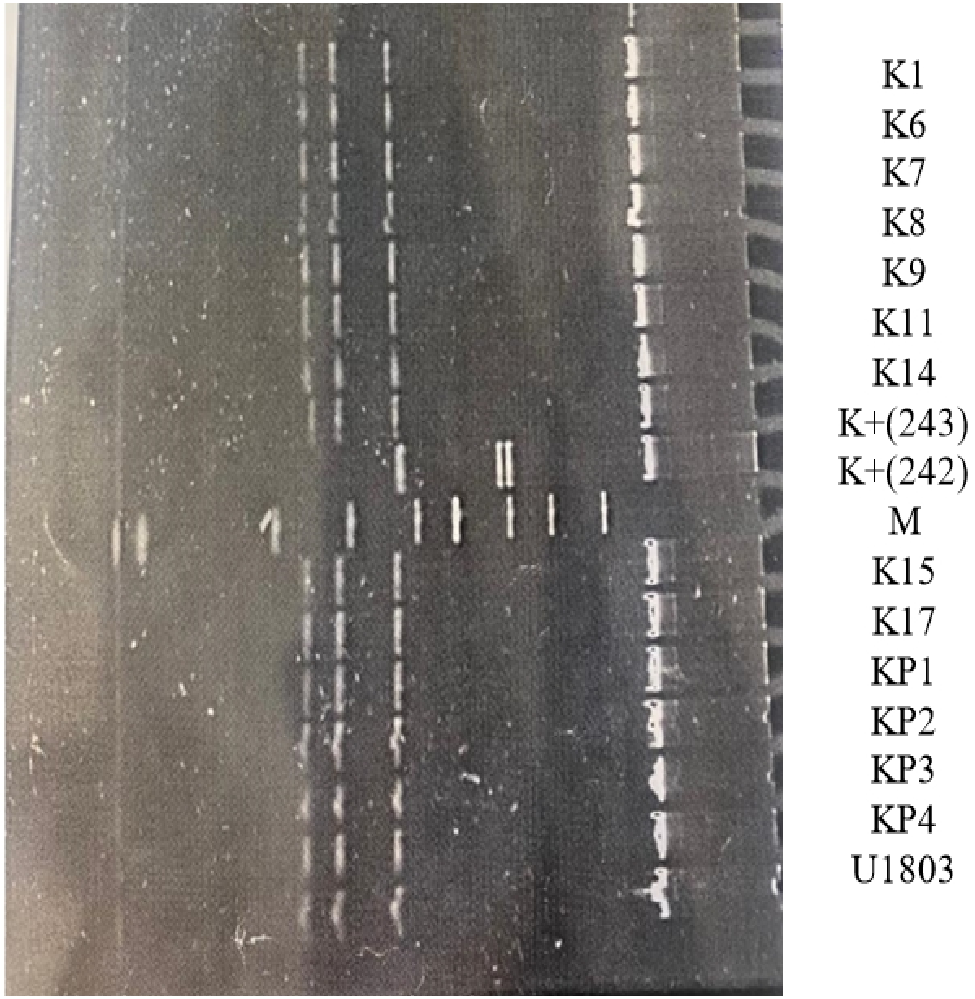
K1
K6
K7
K8
K9
K11
K14
K+(243)
K+(242)
M
K15
K17
KP1
KP2
KP3
KP4
U1803

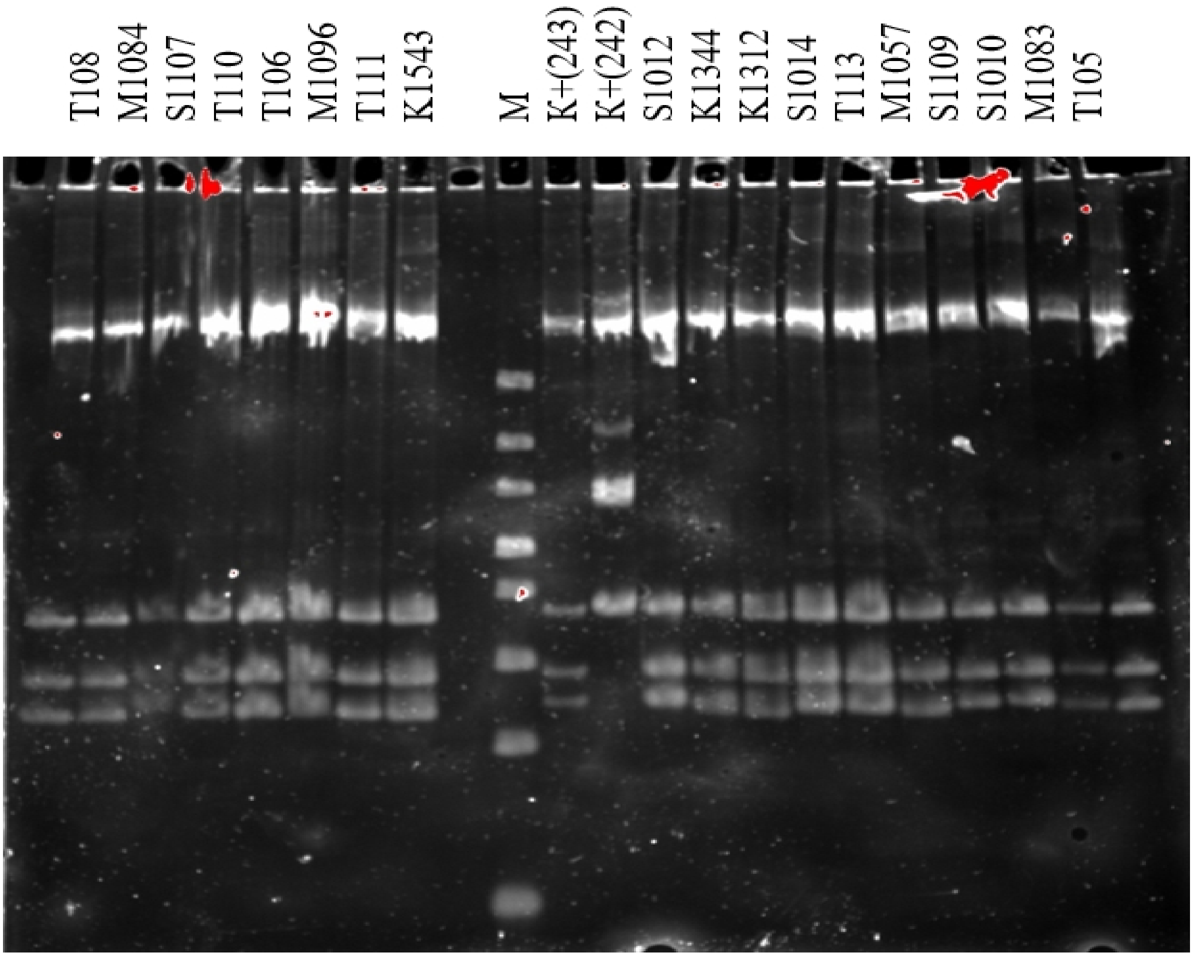
T108
M1084
S1107
T110
T106
M1096
T111
K1543
M
K+(243)
K+(242)
S1012
K1344
K1312
S1014
T113
M1057
S1109
S1010
M1083
T105
