## Supplementary figures and images for "Genetic investigation of honeybee populations in Kazakhstan"

### Figure S2.pdf

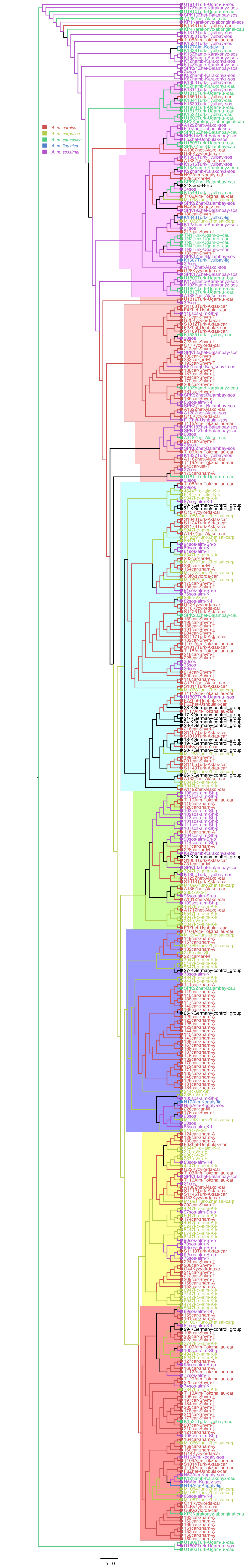
